# Mitochondrial turnover at central GABAergic synapses governs vulnerability to epileptic seizures

**DOI:** 10.64898/2026.06.24.734182

**Authors:** Kristiano Ndoci, Abdulla Chihab, Kiruthika Ganesan, Milica Jevtic, Kelvin M. Tofan, Gulzar Wani, Patric Pelzer, Jorge Soriano-Campos, Franka Odenthal, Vignesh Sakthivelu, Camilla A. Franchino, Laura Pérez-Revuelta, Jonas Benz, Anna-Lena Schumacher, Felix Gaedke, Astrid Schauss, Elisa Motori, Simon E. Troeder, Branko Zevnik, Stefan Mueller, Max Anstötz, Dirk Isbrandt, Matteo Bergami

**Affiliations:** Cologne Excellence Cluster on Cellular Stress Responses in Aging-Associated Diseases (CECAD), University of Cologne, D-50931 Cologne, Germany; Institute of Anatomy II, Medical Faculty, University Hospital Düsseldorf, Heinrich Heine University, 40225 Düsseldorf, Germany; Institute of Biochemistry, University of Cologne, D-50674 Cologne, Germany; Center for Molecular Medicine, 50931 Cologne, Germany; Max Planck Institute for Biology of Ageing, 50931 Cologne, Germany; in vivo Research Facility, Faculty of Medicine and University Hospital Cologne, University of Cologne, Cologne, Germany; Institute of Molecular and Behavioral Neuroscience, Faculty of Medicine, University of Cologne, Cologne 50937, Germany; German Center for Neurodegenerative Diseases (DZNE), Bonn 53175, Germany; University Hospital Cologne, D-50937 Cologne, Germany; Institute of Genetics, University of Cologne, D-50674, Germany

## Abstract

Mitochondrial dysfunction has long been known to underlie neurodegeneration, yet the contribution of mitochondrial turnover dynamics to functional aspects of defined synaptic circuits remains poorly understood. Here, we show that the mitochondrial proteome and turnover rates of hippocampal glutamatergic and GABAergic neurons is differentially remodelled by experience, with distal axon terminals of somatostatin-positive neurons exhibiting most dramatic changes, suggesting a form of metabolic plasticity at GABAergic synapses coupled to circuit activity. Conditional ablation of the mitochondrial transport proteins MIRO1 or TRAK1 (whose human mutations cause congenital epilepsy) stalled turnover at axon terminals driving loss of cristae, without affecting synapse or neuron integrity. The resulting reduction in synaptic GABA levels destabilized network oscillations and led to hyperexcitability, culminating in recurrent seizures and premature death. Post-weaning gene therapy efficiently reversed mitochondrial alterations and ameliorated the epileptic phenotype, underscoring the crucial role of mitochondrial turnover at central GABAergic synapses for balancing network excitability.

## Introduction

Neurons share disproportionate metabolic needs compared to most other cell types on account of their considerably ramified neurites bearing thousands of synapses, whose functionality must be ensured across the entire lifespan. Most of the exceptionally high energetic expenditure faced by neurons is required for maintaining their membrane electrochemical gradients and fire action potentials during synaptic transmission, which are processes largely fuelled by mitochondrial respiration and oxidative phosphorylation (OxPhos) (Harris et al., 2012). These energetic needs are likely further increased in many types of inhibitory GABAergic neurons, which are vastly outnumbered by their glutamatergic counterpart but fire at comparatively much higher frequencies. Within the hippocampus alone, GABAergic neurons exhibit significant phenotypic diversity (Klausberger and Somogyi, 2008), yet they are usually characterized by complex axonal arborisations that require an equally elaborated network of mitochondria localized at their synapses (Kontou et al., 2021). Supporting their naturally elevated metabolic rates (Duarte and Gruetter, 2013), several classes of GABAergic neurons are characterized by a strong immunoreactivity for cytochrome c and cytochrome c oxidase (Gulyas et al., 2006; Kageyama and Wong-Riley, 1982), which are essential for OxPhos, and show a heightened glucose uptake compared to nearby glutamatergic principal neurons (McCasland and Hibbard, 1997). At their presynaptic sites, the extent of cytochrome c labelling correlates with GABAergic neuron activity rates, with terminals of fast-spiking neurons exhibiting denser and more lamellar mitochondrial cristae (Cserep et al., 2018), which are established ultrastructural hallmarks of an enhanced respiratory efficiency (Cogliati et al., 2013). While these studies provide evidence for a metabolic specialization of certain subtypes of GABAergic neurons towards a strong reliance on OxPhos, whether any of these features may be regulated by an animal’s experience – and thus circuit activity – remains unexplored.

Functionally, GABAergic neurons play crucial roles in controlling circuit excitability and coordinating so-called network oscillations (Freund and Buzsaki, 1996). Notably, the energetic costs of network oscillations increase for higher frequency ranges (Kann et al., 2014), and is sustained by mitochondrial OxPhos capacity (Inan et al., 2016; Kann et al., 2011; Whittaker et al., 2011). Network oscillations enable the synchronous discharge of defined ensembles of glutamatergic principal neurons, a process well documented in many brain regions and essential for circuit information processing, memory formation and other higher functions (Buzsaki and Draguhn, 2004). Besides controlling network excitability, many classes of GABAergic neurons also regulate local microcirculation via the activity-dependent release of vasoactive substances (Cauli and Hamel, 2010), which collectively grant these neurons a central role in brain energy metabolism (Buzsaki et al., 2007). The distinctive bioenergetic profile of GABAergic neurons is thought to play important roles in enabling neuronal circuit function (Inan et al., 2016; Olkhova et al., 2023), and even contribute to shape their phenotypic diversity (Moissidis et al., 2025). At the same time, their elevated metabolic demand may lead to higher oxidative stress (Cabungcal et al., 2013), with potential implications for epilepsy and other neurodevelopmental disorders (Marin, 2012). However, which other aspects of the mitochondrial network besides bioenergetics may confer specific vulnerabilities in GABAergic neurons is poorly understood.

Maintenance of functional mitochondria in neurons requires continuous organelle turnover that depends on several intertwined biosynthetic and quality control processes (Basak and Holzbaur, 2025; Misgeld and Schwarz, 2017). The collectively so-called mitochondrial dynamics (anterograde/retrograde motility, immobilization and fission-fusion) are crucial in these processes, ensuring long-term functional integrity of mitochondria throughout complex dendritic and axonal arborizations (Cawley and Farris, 2026; Devine and Kittler, 2018), yet our understanding of how these dynamics may be coordinated in a cell type-specific manner, or even between distinct cellular compartments, is still rudimentary. For instance, the protein Syntaphilin (SNPH) is best characterized for its role in mitochondrial anchoring along axons (Kang et al., 2008), which has direct implications for efficiency of synaptic transmission (Sun et al., 2013), while long-range movement of mitochondria is guaranteed by a molecular complex formed by the integral outer mitochondrial membrane GTPase MIRO1 and the adaptors TRAK1 and TRAK2 (Milton in *Drosophila*), which then bind to dynein and kinesin-1 family motor proteins (Brickley and Stephenson, 2011; Glater et al., 2006; Guo et al., 2005; Macaskill et al., 2009; Stowers et al., 2002; van Spronsen et al., 2013; Wang and Schwarz, 2009). The regulated activity of this machinery allows the positioning and transport of mitochondria to/from dendrites and axons, conceivably ensuring metabolite supply, Ca^2+^ handling and organelle turnover at synaptic sites. However, direct *in vivo* evidence for constitutive mitochondrial turnover at distal terminals has remained elusive until now (Marahori et al., 2024). Moreover, *in vivo* knowledge about mitochondrial motility and distribution processes at central mammalian synapses has mostly focused on glutamatergic neurons (Courchet et al., 2013; Faits et al., 2016; Lewis et al., 2016; López-Doménech et al., 2016; Silva et al., 2021; Smit-Rigter et al., 2016; Sorbara et al., 2014; Takihara et al., 2015), leaving the phenotypically diverse population of GABAergic cells largely unexplored.

Here, we show that experience modifies the neuronal mitochondrial proteome and turnover rates in a cell type– and compartment-specific manner *in vivo*. Using genetic sensors and novel mouse models, we identify a role for axonal mitochondrial transport in enabling distinctively elevated turnover rates at the presynaptic sites of somatostatin (SST)-expressing GABAergic neurons, and show that disturbance of this process via conditional deletion of *Miro1* or *Trak1*, but not *Trak2*, rapidly leads to accumulation of dysfunctional mitochondria in axon terminals, despite lack of neurodegeneration. The ensuing reduction of hippocampal GABAergic neurotransmission drives dysregulated theta/gamma rhythms and network hyperexcitability, and precipitates recurrent seizures leading to increased mortality, thus recapitulating key clinical features linked to *TRAK1* pathogenic variants in human patients.

## Results

### Experience rewires the mitochondrial proteome of GABAergic neurons

To begin investigating the effects of experience on neuronal mitochondrial function, we examined by confocal microscopy the expression levels of two key OxPhos subunits in the mouse hippocampus. We focused on the respiratory chain complex IV (cytochrome C oxidase subunit I, MT-CO1) and on one catalytic portion of the ATP synthase (ATP5B), which we used as proxies to cover potential differences in mitochondrial– and nuclear-encoded OxPhos subunits, respectively. Across all examined hippocampal fields of standard housed (SH) control mice, the CA3 showed the highest expression of both ATP5B and MT-CO1 (Figures 1A and S1A), in line with previous findings showing the highest complex I levels and oxygen consumption rates in this region (Kann et al., 2011). In the remaining major hippocampal sub-fields including the CA1 and dentate gyrus (DG), marker immunoreactivity appeared non-uniform across layers and cell types (Figures 1A and S1A). In these regions, cell somata scattered across the *stratum oriens* and *stratum pyramidale* (in CA1) as well as hilus and inner granule cells layer (in the DG) exhibited most pronounced expression levels, leaving the vast majority of remaining glutamatergic cells located in the principal cell layers (CA1 pyramidal and DG granule neurons) with visibly lower marker expression levels (Figures 1A-1B and S1A). Cells with the highest OxPhos subunit expression within the CA1 and DG sub-fields were immunoreactive for parvalbumin (PV+) or somatostatin (SST+), identifying them as GABAergic neurons (Figures 1B and S1B), supporting previous work (Gulyas et al., 2006). To quantify these differences, we systematically analysed the ratio between ATP5B (or MT-CO1) and TOMM20 (an outer mitochondrial membrane translocase subunit, used here as a normalization marker for mitochondrial mass) at the single-cell level, focussing on PV+, SST+ and glutamatergic neurons in the same brain sections (Figure 1B and S1B). Besides expected variability within even the same class of neuron, analysis confirmed an overall higher relative expression of ATP5B and MT-CO1 in GABAergic cells as compared to glutamatergic ones within the CA1 and DG (i.e., PV– and SST-negative neurons located in the pyramidal and granule cell layers), while this difference evened out in CA3 (Figure 1C). This peculiar pattern in ATP5B and MT-CO1 expression is suggestive of region– and cell type-specific differences that may mirror inherently distinct metabolic profiles between neuron subtypes (Kann et al., 2014). Alternatively, it may reflect the capability of certain cell types to better tune their expression of mitochondrial OxPhos subunits to match changes in microcircuit activity states.

**Figure 1.**
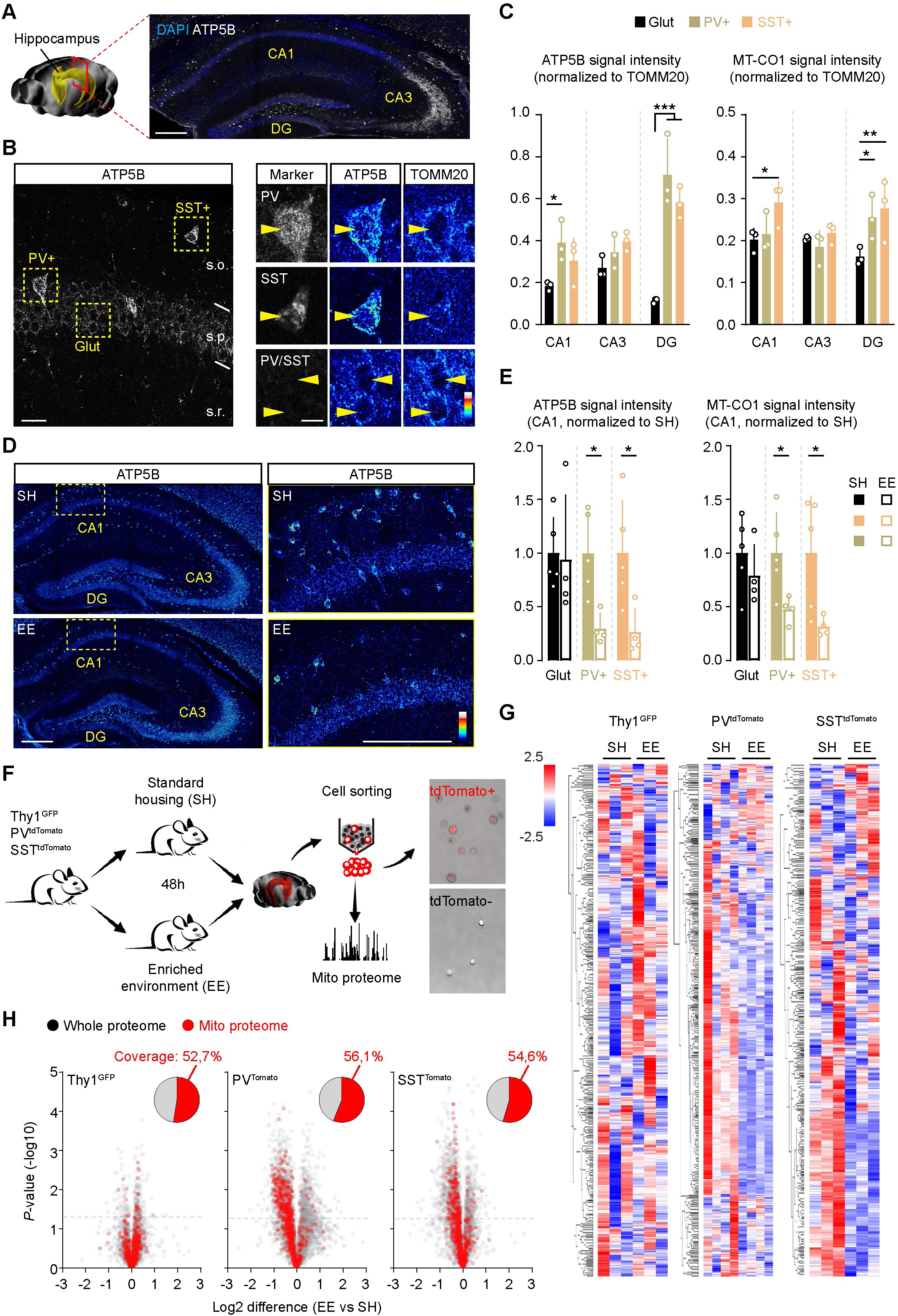
Hippocampal neuron subtype-specific rewiring of the mitochondrial proteome by experience. **(A)** Example of ATP5B immunostaining across hippocampal subfields. Bar, 100 µm. **(B)** Example of ATP5B expression levels in PV+, SST+ and glutamatergic principal neurons in CA1. Right panels depict individual pseudocolored channels for ATP5B and TOMM20. Bars, 15 and 8 µm. **(C)** Quantification of ATP5B and MT-CO1 normalized signal intensity (over TOMM20) for glutamatergic, PV+ and SST+ neurons across hippocampal subfields (n= 3 mice per condition; two-way ANOVA followed by Sidak’s test). **(D)** Hippocampal overviews and zooms showing ATP5B expression levels in mice maintained in SH or exposed to EE for 48h. Bars, 100 µm. **(E)** Quantification of the changes in ATP5B and MT-CO1 signal intensity for CA1 glutamatergic, PV+ and SST+ neurons after EE exposure (n= 4-5 mice per condition; unpaired Mann-Whitney test). **(F)** Experimental design used for assessing proteomic changes induced by EE in hippocampal neurons following cell sorting by flow cytometry of glutamatergic (Thy1-GFP mice), PV+ (PV^tdTomato^ mice) and SST+ neurons (SST^tdTomato^ mice). Right panels show an example of post-sorting confirmation of cell identity. **(G)** Heat-map cluster analysis of the mitochondrial proteome in glutamatergic, PV+, and SST+ neurons of mice maintained in SH or exposed to EE for 48h (n=3-4 mice per condition). **(H)** Volcano plots showing the changes in the mitochondrial proteome (in red; compared to whole proteome, in grey) of EE vs SH mice for glutamatergic, PV+, and SST+ neurons. Cut-off line set at –log10 (*P*-value) = 1.3. Inset shows the mito-proteome coverage according to MitoCarta 3.0. (n=3-4 mice per condition). *, p < 0.05, **, p < 0.01, ***, p < 0.001. See also Figure S1.

To test the latter possibility, we exposed SH adult SST^tdTomato^ reporter mice (SST^Cre^ x tdTomato floxed-stop)(Madisen et al., 2010; Taniguchi et al., 2011) to a novel enriched environment (EE), an established paradigm of increased hippocampal excitability and plasticity (van Praag et al., 2000), and examined the expression levels of ATP5B and MT-CO1 in tdTomato+, PV+ and glutamatergic neurons 2 days later. We observed region-specific changes in the overall expression levels of ATP5B and MT-CO1, with PV+ and tdTomato+ SST neurons within the CA1 and DG exhibiting a significant reduction of signal intensity in enriched versus SH mice (Figures 1D-1E and S1C-S1D), suggesting cell type-specific changes in OxPhos subunit expression levels driven by experience. To broaden our analysis, we performed an unbiased characterization of the changes in somatic mitochondrial proteome induced by EE of all 3 examined hippocampal neuron classes via cell sorting by flow cytometry followed by label-free proteomics (Figure 1F). Cell sorting was performed on rapidly dissociated hippocampi obtained from 3 validated mouse lines bearing each a fluorescent reporter restricted to the respective neuronal population of interest: Thy1-GFP for glutamatergic neurons (Feng et al., 2000); PV^tdTomato^ (PV^Cre^ x tdTomato floxed-stop mice) for PV+ neurons (Hippenmeyer et al., 2005); and SST^tdTomato^ mice. After visual confirmation of cell type-specific enrichment (Figure 1F), mass spectrometry analysis revealed a comparable mitochondrial proteome coverage between mouse lines (Thy1-GFP: 52,7%; PV^tdTomato^: 56,1%; and SST^tdTomato^: 54,6%) according to MitoCarta 3.0 (Rath et al., 2021). This allowed us to confirm that the expression levels of many OxPhos subunits across virtually all respiratory complexes was indeed enhanced specifically in PV+ and SS+ neurons compared to glutamatergic ones in SH (Figure S1E). Hierarchical clustering of this dataset further showed that cells sorted from PV^tdTomato^ and SST^tdTomato^ mice underwent most prominent changes of their somatic mitochondrial proteome following EE (Figure 1G). Besides the expected variability between samples of even the same mouse line, GABAergic neurons exhibited a noticeable fraction of mitochondrial proteins undergoing significant changes in their relative abundance, mostly towards a downregulation, with PV+ neurons showing a greater effect (29.3% of detected mitochondrial proteins) compared to SST+ neurons (6.9%, *q* value < 0.05) (Figure 1H). In contrast, glutamatergic neurons showed minimal changes in their mitochondrial proteome (Figure 1H). These dynamics were mirrored by a net negative shift in the relative abundance of the mitochondrial proteome as a whole compared to the rest of the cell proteome of SST+ and PV+ neurons (Figure S1F-S1G), despite no visible reduction in somatic mitochondrial mass (Figure S1H). Accordingly, KEGG pathway analysis of SST+ and PV+ neurons disclosed a significant down-regulation of numerous metabolic and mitochondrial processes (among others), including amino acid and lipid metabolism, as well OxPhos (Figure S1I).

Together, these data indicate that a relatively short period of experience elicits a distinctive rewiring of the mitochondrial proteome in PV+ and SST+ GABAergic neurons, presumably as an adaptive mechanism to cope with changes in circuit activity and metabolic needs.

### Neuron subtype-specific differences in mitochondrial turnover induced by experience

The observation that a short (48h) EE exposure preferentially drives mito-proteome changes in PV+ and SST+ GABAergic neurons over glutamatergic ones, prompted us to broaden our investigation to other aspects of mitochondrial plasticity that may occur during a similar time scale. Mitochondrial turnover is widely accepted as an important form of quality control critical for maintenance of neuronal survival (Misgeld and Schwarz, 2017), but neuronal subtype-specific differences in turnover have been challenging to report. To capture relative changes in mitochondrial turnover rates *in vivo*, we generated a Cre-dependent adeno-associated viral vector (AAV) encoding for the previously optimized, doxycycline (Dox)-dependent genetic sensor mitoTimer (Hernandez et al., 2013). Upon expression, mitoTimer acts as a molecular clock by shifting its emission spectra from green to red over the course of 48h, with any additional shot of Dox resulting in new green fluorophore influx into the existing pool of previously matured mitochondria, allowing to timely quantify relative changes in organelle turnover by ratioing green over red signals (Figure 2A). We tested mitoTimer kinetics in neuronal cultures (Figures S2A-S2B) and *in vivo*, targeting either glutamatergic pyramidal neurons within the somatosensory cortex or granule neurons in the hippocampal DG (Figures S2C-S2F). In both settings, increasing the frequency of Dox shots after an initial application 3 days earlier had cumulative effects, resulting in the progressive shift of the signal ratio towards a “greener” mitoTimer range, particularly in the somato-dendritic compartment, where mitochondrial biogenesis and fusion dynamics are expected to be predominant. Given that mitochondrial biogenesis, degradation and transport are all cellular processes influencing mitochondrial turnover rates as reported by mitoTimer (Ferree et al., 2013; Hernandez et al., 2013), we reasoned that comparing putative changes between soma and axon terminals of the same neuron type would reflect the prevalence of compartment-specific processes (that is, higher biogenesis within the soma versus a more pronounced role of transport for axon terminals) (Pastor et al., 2025). In addition to hippocampal DG granule neurons, we therefore targeted mitoTimer selectively to PV+, SST+ and glutamatergic pyramidal cells in CA1 taking advantage of mouse and viral CRE-dependent recombination approaches (Figure 2B). To synchronize new mitoTimer expression across cell types, we delivered a first Dox pulse and waited 3 days to allow complete fluorophore maturation, followed by imaging at 1 day after a final Dox pulse in presence/absence of EE (Figure 2B), which is a time interval compatible with the previously observed changes in mitochondrial proteome (Figures 1F). We then examined the green/red signal ratio in the targeted neuron types (Figure 2C), specifically comparing their soma versus axon terminals. Imaging analysis revealed a broad distribution in terms of green/red signal ratios across mitochondria within neuronal compartments of even the same cell type (Figure 2D), making it difficult to compare cell type-specific differences simply averaging whole fields of view. Instead, we generated masks for individual mitochondrial region of interest (ROI) within each images (e.g., neuronal cell body), and quantified for each ROI its own signal ratio, arbitrarily establishing threshold values below and above which we defined mitochondria for having overtly lower (redder, likely reflecting the accumulating of “older” mitoTimer) or higher (greener, reflecting increased new mitoTimer incorporation) turnover rates compared to the bulk of intermediate ratios (Figure S2G). Specific analysis of the proportion of mitochondria falling within each of these two categories revealed clear differences between somatic regions of glutamatergic (CA1 pyramidal and DG granule cells) and GABAergic neurons (SST+ and PV+ in CA1) after EE, with the former accumulating redder mitochondria and thus exhibiting overall lower turnover rates than GABAergic cells (Figures 2D, 2E and S2H). In contrast, the proportion of greener mitochondria within the soma appeared similar across cell types, suggesting uniform rates of biogenesis following 2 days of EE (Figure 2E).

**Figure 2.**
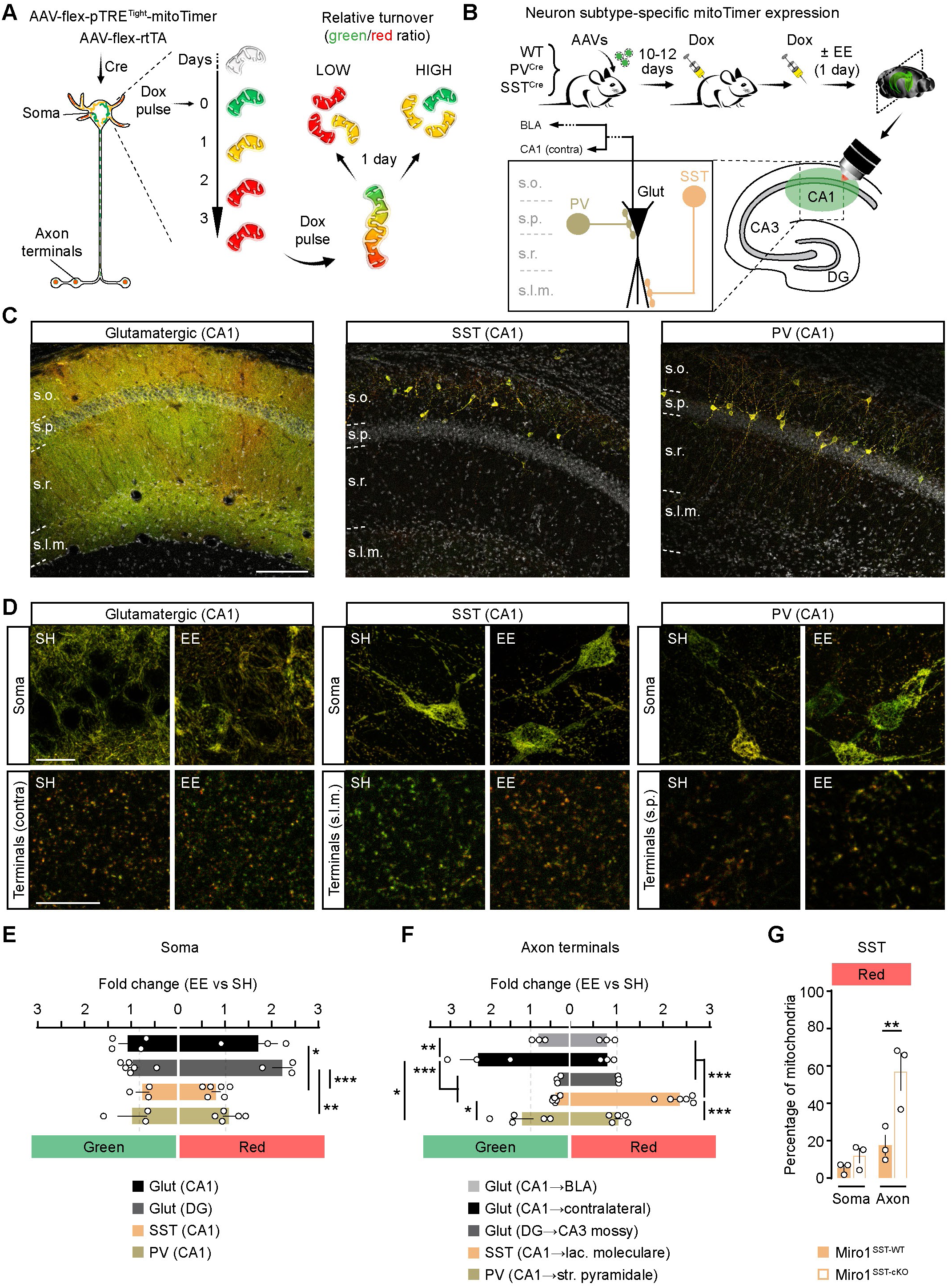
Experience differentially modulates mitochondrial turnover of glutamatergic and GABAergic neurons at their soma and axon terminals. **(A)** Scheme depicting the mode-of-action of the used Cre– and Dox-dependent AAV expressing mitoTimer to target hippocampal neurons *in vivo*. Three days after an initial pulse of Dox, most mitochondria have turned red. A second pulse of Dox under these settings induces new mitoTimer production, enabling the ratiometric analysis of cell compartment-specific changes (high vs low relative turnover) after a day. **(B)** Experimental plan used to target mitoTimer to glutamatergic, PV+ and SST+ neurons within the CA1 *in vivo*, followed SH/EE exposure and imaging analysis. Paradigms involved the use of Cre-expressing lines target PV+ and SST+ neurons, or the co-injection of a CAMKII-Cre expressing AAV to target glutamatergic neurons. Inset shows the anatomical organization of the targeted neurons and their analyzed somatic/axonal compartments. **(C)** Overviews of the CA1 region showing mitoTimer expression (merged green and red signals) in the targeted neuronal populations. Bar, 70 µm. **(D)** Representative zooms of glutamatergic, SST+ and PV+ neurons (top, soma; bottom, axonal terminals at the indicated locations) in mice exposed to EE or SH. Bar, 20 µm. **(E)** Quantification of mitoTimer signal ratio after EE within the cell body and **(F)** axon terminals of the indicated neuronal populations (n= 3-6 mice per condition; one-way ANOVA followed by Tukey’s test). **(G)** Quantification of “high red” mitoTimer signal within soma and axon terminals of Miro1^SST-WT^ and Miro1^SST-cKO^ mice (n= 3 mice per condition; one-way ANOVA followed by Tukey’s test). *, p < 0.05, **, p < 0.01, ***, p < 0.001. S.o., stratum oriens; s.p., stratum pyramidale; s.r., stratum radiatum; s.l.m., stratum lacunosum-moleculare. See also Figure S2.

Unexpectedly, mitochondrial turnover at the axon terminals of most cell types did not match its corresponding pattern at their soma. Long-range terminals of CA1 pyramidal neurons projecting to the contralateral hippocampus or to the basolateral amygdala (BLA), as well as mossy fibre terminals of DG granule neurons in CA3, appeared visibly redder than their somatic counterpart already in SH, and EE exposure left them marginally changed within the red range (Figures 2D, 2F, S2G and S2H). A similar pattern was observed for PV+ neurons, whose axon (for so-called basket cells) ramifies locally to densely innervate the soma of nearby pyramidal cells (Freund and Buzsaki, 1996). Interestingly, there was a significant difference in the rather small mitochondrial high-green fraction (Figure S2G) between distinct axonal projections of glutamatergic neurons, with mitochondria in contralateral CA1 terminals undergoing more apparent turnover than those in BLA terminals, or even of CA3 mossy fibre terminals (Figure 2F), possibly reflecting projection-specific differences in mitochondrial transport or local degradation. By contrast, the terminals of SST+ neurons (so-called *oriens/lacunosum-moleculare* cells, or OLM) whose axon extends into the *stratum lacunosum-moleculare* to profusely innervate the distal apical tuft dendrites of CA1 pyramidal cells (Freund and Buzsaki, 1996), contained noticeably greener mitochondria than those of other cell types in SH, and their signal sharply shifted towards red in response to EE (Figures 2D and 2F), consistent with a reduction in apparent turnover rates at this site. To experimentally assess the contribution of transport to the distinguishing axonal mitochondrial turnover rates of SST+ neurons, we performed the same experiment after conditional deletion of the outer mitochondrial membrane GTPase MIRO1, which is required for axonal mitochondrial trafficking to synapses (Guo et al., 2005; Macaskill et al., 2009), by crossing *Miro1* floxed (Nguyen et al., 2014) with SST^Cre^ mice. MitoTimer analysis revealed a prominent accumulation of redder mitochondria at the axon terminals of MIRO1-deficient SST+ neurons, while their soma exhibited comparatively more modest changes (Figures 2G and S2I), revealing a substantial role for mitochondrial transport in maintaining basal turnover rates at their distal sites.

These data support the existence of experience-driven, and cell type-specific differences in relative mitochondrial turnover across neuronal compartments, with SST+ neurons showing pronounced turnover rates at their terminals already at basal levels, which can be blocked by interfering with mitochondrial transport dynamics.

### Conditional disruption of mitochondrial transport in GABAergic neurons

To understand the relevance of distal mitochondrial turnover dynamics for GABAergic neuron synapse function, we next sought to genetically interfere with the molecular machinery regulating long-range mitochondrial transport, starting with MIRO1 and extending the analysis to the motor adaptor proteins TRAK1 and TRAK2. We first took advantage of available *Miro1* floxed animals to validate in neuronal culture the requirement of MIRO1 for axonal transport by performing loss of function and rescue experiments via AAV-based expression of CRE recombinase and wild-type MIRO1 (Figures S3A-S3E). Through crossings with SST^Cre^, PV^Cre^ or (pan-GABAergic) Vgat^Cre^ mice (Vong et al., 2011), we then generated mouse lines ensuring the conditional deletion of *Miro1* specifically in SST+ (Miro1^SST-cKO^ mice), PV+ (Miro1^PV-cKO^ mice) or the whole population of GABAergic neurons (Miro1^Vgat-cKO^ mice). Our crosses also included reporter lines for either cytosolic tdTomato, mitochondrial-targeted YFP (mtYFP), or both, to monitor gene recombination rates, cell fate, and putative mitochondrial phenotypes in the resulting mice. All lines were born at mendelian ratios, indicating that even prenatal deletion of *Miro1* via the Vgat^Cre^ line (recombining earlier compared to SST^Cre^ and PV^Cre^ line used)(Hippenmeyer et al., 2005; Taniguchi et al., 2011; Vong et al., 2011) did not majorly impact embryonic development. This was further corroborated by analysis of body weight and brain-specific structural parameters such as overall architecture, cortex area, and number/distribution of recombined GABAergic neurons in both somatosensory cortex and hippocampal CA1 of Miro1^Vgat-cKO^ pups at birth, which were similar to control littermates (Figures S3F-S3H). However, all Miro1^Vgat-cKO^ pups showed perinatal lethality within the first 24h, which was preceded by acquisition of a cyanotic aspect despite their virtually unremarkable behaviour at birth, indicating that ablation of MIRO1 in the vast majority of GABAergic neurons is not compatible with postnatal life, a phenotype similar to that of *Miro1* knock-out mice (Nguyen et al., 2014). In contrast to Miro1^Vgat-cKO^ mice, Miro1^SST-cKO^ and Miro1^PV-cKO^ mice survived well into adulthood, and their growth as well as density of recombined tdTomato+ neurons across neocortex and hippocampus were virtually indistinguishable from those of littermate controls at the examined ages (up to 2 months for Miro1^SST-cKO^ and 12 months for Miro1^PV-cKO^ mice) (Figures 3A-3B, S3I-S3J, and S3M-S3O), despite validation of MIRO1 loss (Figure S3K). Single cell analysis of SST+ and PV+ neurons in hippocampal CA1 of control mice revealed a uniformly distributed mitochondrial network across soma and dendrites, with a progressive age-dependent shift from small to more tubular or even visibly elongated morphologies in dendrites over the course of 8 weeks (Figures 3C, S3L and S3P), in line with similar reports in other types of mouse neurons *in vivo* (Faitg et al., 2021; Kochan et al., 2024; Lewis et al., 2018; Popov et al., 2005; Virga et al., 2024). Starting from 4 weeks, deletion of *Miro1* caused a moderate mitochondrial fragmentation within the cell soma (Figure S3L), which progressed to more severe changes by 2 months of age, leaving mitochondria with fragmented morphologies across the whole somato-dendritic compartment, in conjunction with the formation of conspicuous clusters (Figures 3C and S3P), in agreement with previous studies (Kontou et al., 2021). Quantification of volumetric mitochondrial occupancy revealed a significant reduction within both soma and dendritic compartments of MIRO1-deficient neurons, while axonal boutons showed no quantifiable changes (Figures 3C-3D and S3P-S3Q), on account of the already much smaller size of mitochondria at axon terminals of control SST+ and PV+ neurons (Figures 3C and S3P), suggesting that no mitochondrial depletion took place at this distal site. To further corroborate this point, we quantified the percentage of axonal boutons containing mitochondria of double-positive (tdTomato/mtYFP) SST+ neurons in the CA1 *lacunosum-moleculare*, and found no differences between genotypes (roughly 80% of boutons contained mitochondria), despite a small but significant reduction of the average mitochondrial volume in Miro1^SST-cKO^ mice (Figures 3E-3F).

**Figure 3.**
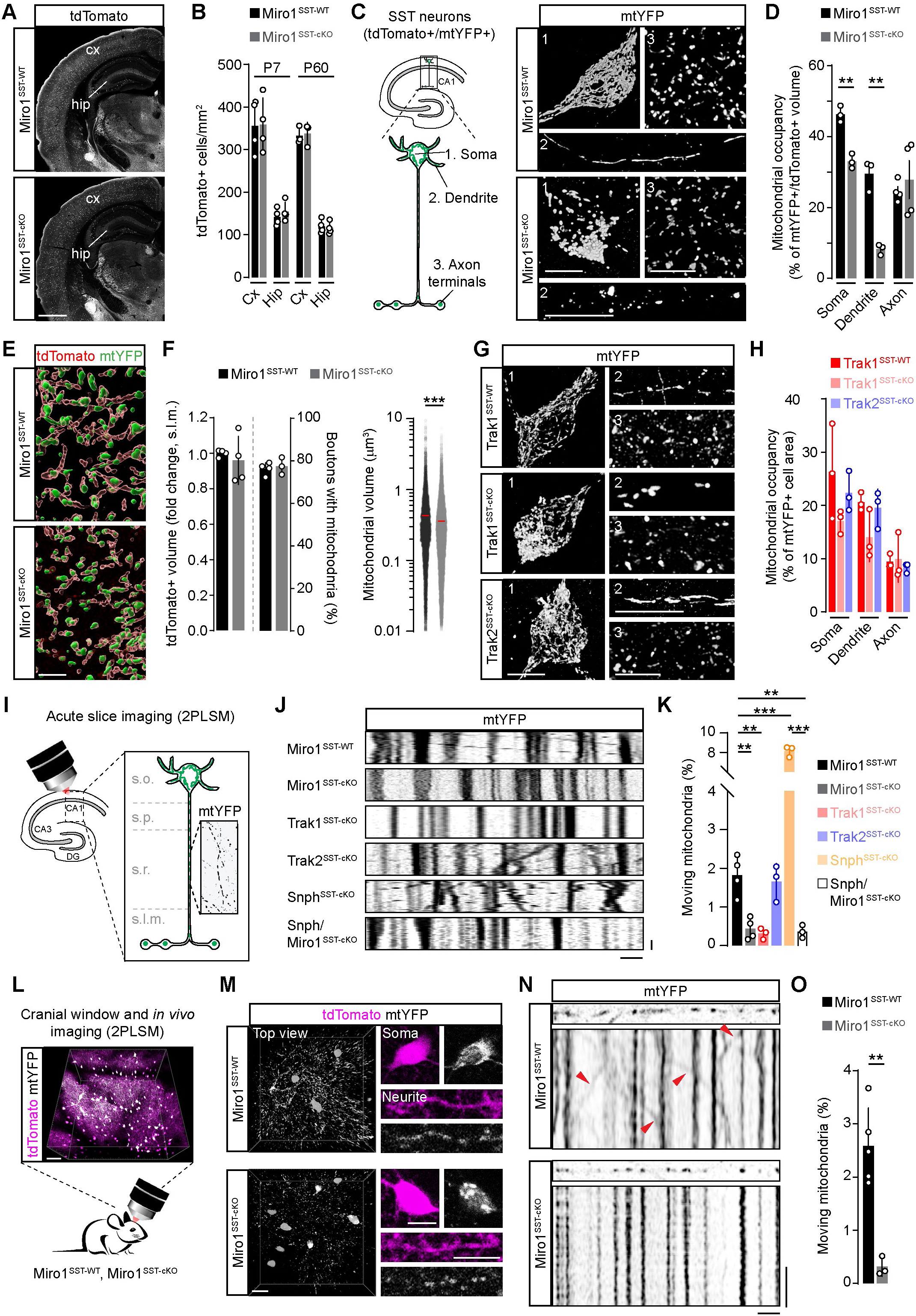
Disrupted mitochondrial morphology and axonal transport in SST+ GABAergic neurons lacking *Miro1* or *Trak1*. **(A)** Examples of adult Miro1^SST-WT^ and Miro1^SST-cKO^ brain sections expressing tdTomato. Bar, 800 µm. **(B)** Quantification of cortical and hippocampal tdTomato+ neurons at P7 and P60 in Miro1^SST-WT^ and Miro1^SST-cKO^ mice (n= 3-4 mice per condition; two-way ANOVA followed by Sidak’s test). **(C)** Scheme depicting the anatomical organization of CA1 SST+ neurons and their analyzed cellular compartments. Pictures on the right illustrate the mitochondrial morphology within soma, dendrites and axon terminals of mtYFP-expressing SST+ neurons in presence and absence of MIRO1. Bars, 8, 8 and 4 µm. **(D)** Quantification of mitochondrial occupancy (over tdTomato+ cell volume) across neuronal compartments in Miro1^SST-WT^ and Miro1^SST-cKO^ mice (n= 3-4 mice per condition; unpaired t-test). **(E)** Volumetric reconstruction of axon terminals in mtYFP/tdTomato expressing CA1 SST neurons of Miro1^SST-WT^ and Miro1^SST-cKO^ mice. Bar, 3 µm. **(F)** Quantification of boutons containing mitochondria (left; n= 3-4 mice per condition; unpaired t-test) and mitochondrial volume in axon terminals (right; n > 10.000 mitochondria per condition; unpaired t-test) of Miro1^SST-WT^ and Miro1^SST-cKO^ mice. **(G)** Examples illustrating the mitochondrial morphology within soma, dendrites and axon terminals of mtYFP-expressing SST+ neurons in presence and absence of TRAK1 or TRAK2. Bars, 8, 8 and 4 µm. **(H)** Quantification of mitochondrial occupancy (over area) across neuronal compartments in Miro1^SST-WT^ and Miro1^SST-cKO^ mice (n= 3 mice per condition; one-way ANOVA followed by Dunnett’s test). **(I)** Scheme depicting the ROI in the CA1 s.r. examined by 2PLSM in acute hippocampal slices to monitor mitochondrial trafficking in mtYFP-expressing SST+ neurons. **(J)** Representative kymographs of the indicated genotypes. Bars, 4 µm and 1 min. **(K)** Quantification of mitochondrial motility in the indicated genotypes (n= 3-4 mice per condition; one-way ANOVA followed by Tukey’s test). **(L)** Scheme depicting the intravital cortical imaging mtYFP/tdTomato-expressing SST+ neurons. Bar, 50 µm. **(M)** Examples of imaged neurons in Miro1^SST-WT^ and Miro1^SST-cKO^ mice showing mitochondrial network in cell body and neurites. Bar, 8 µm. **(N)** Representative *in vivo* kymographs of SST+ neurons in Miro1^SST-WT^ and Miro1^SST-cKO^ mice. Arrowheads point to moving mitochondria. Bars, 4 µm and 1 min. **(O)** Quantification of mitochondrial motility as shown in N (n= 3-5 mice per group; Welch’s t-test). *, p < 0.05, **, p < 0.01, ***, p < 0.001. See also Figure S3.

We next asked whether these alterations in GABAergic neuron mitochondrial network were caused by a disruption of transport, or were rather specific consequences of MIRO1 loss, as this protein has been credited several additional roles in mitochondrial structural and quality control processes besides transport itself (Konig et al., 2021; López-Doménech et al., 2021; Modi et al., 2019; Nemani et al., 2018; Saotome et al., 2008; Shlevkov et al., 2016). Instead of examining each of these proposed roles separately, we generated floxed animals for *Trak1* (exon 5) and *Trak2* (exon 4) genes (Figure S4A), to independently target mitochondrial transport along microtubules in axons and dendrites, and performed a similar set of analyses while preserving the expression of MIRO1. Validation of *Trak1* and *Trak2* conditional deletion was first conducted in neuronal cultures, where transduction with a CRE-expressing AAV confirmed protein loss in both models via western blot analysis (Figure S4B). Likewise, kymograph studies of CRE-GFP and GFP-only (control) expressing neurons co-labelled with MitoTraker Red showed a significant reduction of mitochondrial motility along axons of TRAK1-deficient cells, and a less prominent but not significant trend in dendrites (Figures S4C-S4D). In contrast, TRAK2-deficient neurons showed a predominant defect in dendritic but not axonal mitochondrial transport (Figures S4E-S4F), which together support earlier knock-down findings in neuronal cultures (Brickley and Stephenson, 2011; van Spronsen et al., 2013). To discern the role of TRAK1 and TRAK2 in GABAergic neurons, we then crossed floxed animals with Vgat^Cre^ and mtYFP reporter mice. Like for Miro1^Vgat-cKO^ mice, pups of both resulting lines were born at mendelian ratios, indicating that knock-out of *Trak1* or *Trak2* is compatible with embryonic development. However, during the first postnatal days virtually all Trak1^Vgat-cKO^ mice developed a stereotyped phenotype consistent with repetitive spasms and twitching movements, which is reminiscent of the congenital hyperekplexia and myoclonus of infant patients carrying *TRAK1* pathogenic variants, who often develop refractory status epilepticus leading to premature death within few years (Barel et al., 2017; Li et al., 2024a; Sagie et al., 2018). By day 7, this phenotype in Trak1^Vgat-cKO^ mice progressed into limb rigidity and tail stiffness (compatible with hypertonia) culminating in pups laying on one side and precluding their ability to properly feed and move (Figure S4G). This led to a significant reduction in weight gain (Figure S4H) followed premature death, which (as for Miro1^Vgat-cKO^ mice) required early culling of animals for ethical reasons. Inspection of Trak1^Vgat-cKO^ brains at P7 revealed a surprisingly overall intact cytoarchitecture and density of recombined GABAergic neurons (within neocortex and hippocampus), emphasizing the lack of overt neurodegeneration in this mouse model (Figures S4I-S4J), despite the presence of visible mitochondrial clusters in a fraction of the cells when examined by immunohistochemistry (Figure S4I). In contrast, Trak2^Vgat-cKO^ mice did not exhibit any visible phenotype, sign of premature death or broad cellular alterations including in somatic mitochondrial morphology up to 1 year of age (Figures S4K-S4M). We next proceeded to examine Trak1^SST-cKO^ and Trak2^SST-cKO^ mice, which predictably grew well into adulthood up to 3-4 months of age, exhibiting intact brain cytoarchitecture and density of recombined neurons (Figure S4N-S4P). By 2 months, however, mtYFP+ neurons in Trak1^SST-cKO^ mice showed a consistent mitochondrial fragmentation phenotype within their somato-dendritic compartment (Figures 3G-3H), which although less pronounced was reminiscent of the one identified in Miro1^SST-cKO^ mice.

To link the alterations in mitochondrial morphology of Miro1^SST-cKO^ and Trak11^SST-cKO^ mice with defects in mitochondrial transport within the native hippocampal tissue, we took advantage of the unique anatomy of CA1 SST+ neurons, whose thin axon cuts across the entire *stratum radiatum* before arborizing, thus enabling the quantification of mitochondrial motility along a largely linear axonal segment in acute explants (Figures 3I). Live imaging by 2-photon laser scanning microscopy (2PLSM) of mtYFP-expressing axons in acute brain slices revealed that about 2% of labelled mitochondria were indeed motile in 3-week-old control neurons during imaging sessions of 3 minutes, yet this percentage dropped by 4-5 folds in Miro1^SST-cKO^ and Trak1^SST-cKO^ neurons, virtually ablating motility, while it was left unaffected in Trak2^SST-cKO^ mice (Figures 3J-3K). In an effort to genetically rescue the motility defect in these mouse models, we generated transgenic animals by engineering an artificial double-floxed stop codon-containing cassette into intron 4 of the mitochondrial docking gene *Syntaphilin* (*Snph* flexed mice), whose CRE-mediated recombination led to increased mitochondrial motility in neuronal cultures (Figures S4Q-S4R) and CA1 SST+ neurons (assessed in Snph^SST-cKO^ mice) by about 4-fold (Figures 3J-3K). However, simultaneous deletion of *Snph* and *Miro1* in the same neurons (Snph/Miro1^SST-cKO^ mice) led to no appreciable recovery of axonal mitochondrial transport, indicating the essential requirement of MIRO1 in this process (Figures 3J-3K). Importantly, we validated *in vivo* our key *ex vivo* findings by performing intravital imaging of anesthetized Miro1^SST-cKO^ and littermate control mice (Figure 3L). Using this imaging setting, and focusing our analysis on accessible layers II/III of the somatosensory cortex, we corroborated (i) the disruption of mitochondrial morphology within soma and neurites of double-positive tdTomato/mtYFP neurons in absence of chemical fixation (Figure 3M), as well as (ii) the impairment in mitochondria motility along neurites after *Miro1* deletion (Figures 3N-3O).

Together, these data demonstrate that while ablation of either *Miro1* or *Trak1* does not drive visible degeneration in GABAergic neurons, it efficiently disrupts their axonal mitochondrial transport, leading to subsequent alterations in organelle morphology. Moreover, *Trak1* deletion phenocopies the mitochondrial defects observed in absence of MIRO1, indicating a primary role of mitochondrial transport for both genes in GABAergic SST+ neurons.

### Ablation of *Miro1* or *Trak1* in SST+ neurons leads to recurrent seizures

By 8 weeks of age, some Miro1^SST-cKO^ mice of both sexes developed sporadic seizures (lasting in average less than 1 minute) (Figure 4A) characterized by involuntary head and forelimb jerking movements. The percentage of mice showing seizures steadily increased in the following weeks, and although animals appeared to fully recover after these initial events, over time this pattern was followed by a corresponding gradual decrease in the survival rate (Figure 4B), with animals found suddenly dead even without prior signs of distress during cage inspection in the same day. At the histological level, the immunoreactivity of markers for astrocytic and microglial gliosis (GFAP and IBA1) was only slightly changed in Miro1^SST-cKO^ mice (Figures S5A-S5C), despite the presence of recurrent seizures, but consistent with lack of overt neurodegeneration. By about 12 weeks, animals began developing visible convulsions consistent with generalized tonic-clonic seizures, characterized by rearing, stiffness, and falling with fore– and hindlimb jerking movements, which could potentially explain their sudden death. Interestingly, no such convulsing phenotype was observed in Miro1^PV-cKO^ mice. Reflecting their impairment in mitochondrial motility and disrupted mitochondrial morphology, Trak1^SST-cKO^ (but not Trak2^SST-cKO^) mice developed a similar phenotype, albeit delayed by several weeks, with a seizure onset around 14 weeks of age then progressing to tonic-clonic convulsions and to an increased mortality rate compared to controls (Figures 4A-4B). Strikingly, Snph/Miro1^SST-cKO^ mice faithfully reproduced, or even slightly worsened, the phenotype of Miro1^SST-cKO^ mice (Figures 4A-4B), consistent with their failure in rescuing motility defects in SST+ neurons.

**Figure 4.**
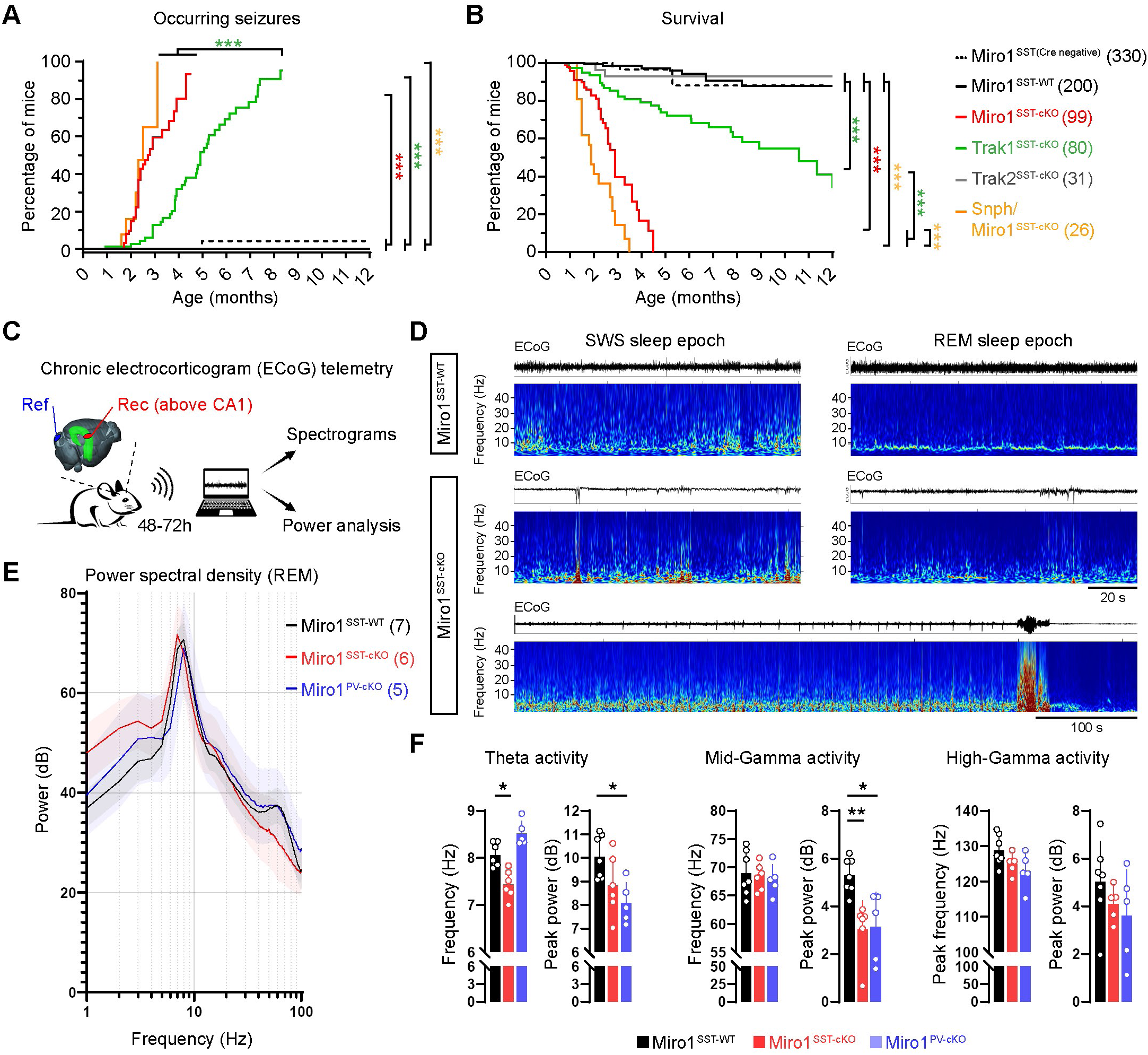
*Miro1* and *Trak1* mutant mice spontaneously develop epileptic seizures. **(A)** Graphs depicting the incidence of seizures and **(B)** survival rate over time in the indicated mouse genotypes (Mantel-Cox test; experimental n written in brackets). **(C)** Scheme of the EcoG setup used, showing the placement of recording (above CA1) and reference (above cerebellum) electrodes. **(D)** Example of ECoG traces (top) and corresponding wavelet spectrograms (bottom) recorded from Miro1^SST-WT^ and Miro1^SST-cKO^ mice during SWS and REM sleep stages. The recoding at the very bottom reports on a seizure event. **(E)** Averaged power spectral analysis during REM sleep for the indicated genotypes. **(F)** Frequency and power quantifications in the theta and gamma range for the indicated genotypes (n= 5-7 mice per group; Kruskal-Wallis followed by Dunn’s test). *, p < 0.05, **, p < 0.01, ***, p < 0.001. See also Figure S5.

To corroborate our visual assessment and quantify potential neuronal network activity alterations in Miro1^SST-cKO^ mice, we performed chronic electrocorticogram (ECoG) recordings in the somato-motor cortical area (above dorsal CA1) over the course of 48-72h (Figure 4C). This allowed us to capture the presence of interictal spikes during both slow-wave (SWS) and REM sleep, and confirm cases of spontaneous seizures in Miro1^SST-cKO^ mice (Figure 4D), but not in Miro1^PV-cKO^ animals (Figure S5D). Yet, despite lack of spontaneous seizures in Miro1^PV-cKO^ mice, power spectral density analysis (Figure 4E) revealed a specific reduction in theta and mid-gamma bands (i.e., frequency and/or power) during REM sleep in both Miro1^PV-cKO^ and Miro1^SST-cKO^ mice (Figure 4F), which would be consistent with the role SST+ and PV+ neurons play in regulating network oscillations at these frequencies (Klausberger and Somogyi, 2008), demonstrating circuit activity abnormalities linked to the corresponding neuronal sub-population targeted.

These data indicate that GABAergic neuron-specific disruption of mitochondrial transport mediated by MIRO1 and TRAK1 alters neuronal network activity, and identifies a crucial role of these genes in preventing the development epileptogenic phenotypes in mice.

### Deletion of *Miro1* in SST+ neurons impairs GABAergic transmission onto nearby glutamatergic neurons

To better ascertain the consequences of mitochondrial transport disruption in SST+ neurons for network excitability, we performed electrophysiological recordings of Miro1^SST-cKO^ mice and control littermates at the presymptomatic age of 5-6 weeks, which precedes the onset of seizures at around 8 weeks. SST+ neurons in the *stratum oriens* of the CA1 region were visually identified in acute hippocampal slices by the expression of the fluorescent reporter tdTomato (Figures 5A-5B), and further characterized during whole-cell patch-clamp recordings by their firing pattern into previously defined “high” and “low” firing neuronal subtypes (Hewitt et al., 2021), of which only the “high” firing group was represented in sufficient numbers in our dataset for subsequent analysis. Recordings revealed no overt alterations in the resting membrane potential, input resistance, action potential threshold, current-voltage relationship, and firing pattern in Miro1^SST-cKO^ mice compared to controls (Figures 5C-5D and S5E-S5F). Moreover, no differences in frequency and amplitude of spontaneous post-synaptic currents (sPSCs) were identified (Figures 5E-5F). Accordingly, analysis of EGFP-Gephyrin and Psd95-GFP expression via AAV-mediated *in vivo* delivery confirmed a very similar density of post-synaptic inhibitory and excitatory foci (both perisomatic and dendritic) between CA1 SST+ neurons of Miro1^SST-cKO^ and control littermates (Figures S5G-S5I), collectively indicating that *Miro1* deletion in hippocampal SST+ neurons does not overtly affect their innervation and synaptic integration into the network

**Figure 5.**
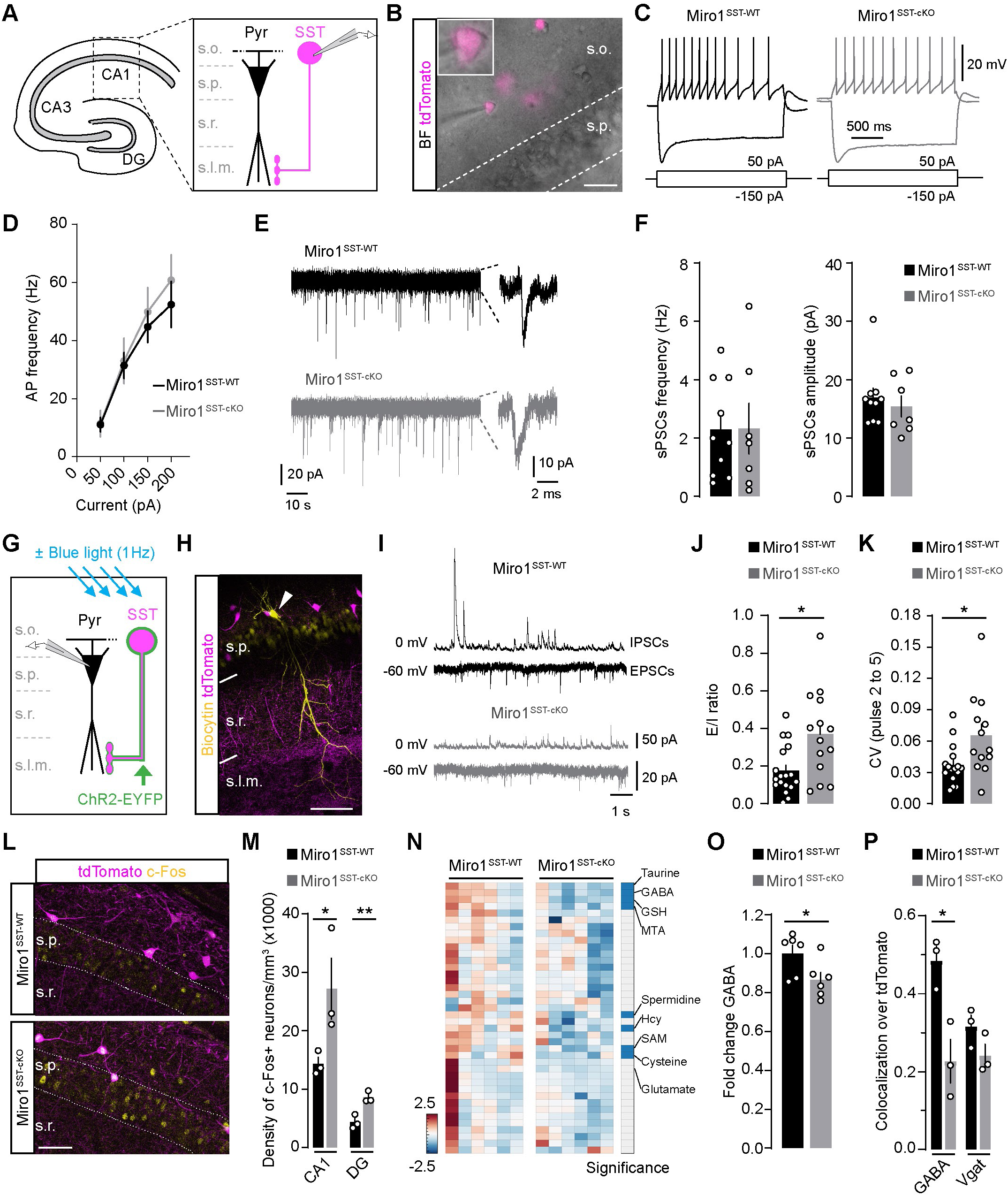
Unchanged membrane properties but reduced GABAergic synaptic transmission of SST neurons lacking *Miro1*. **(A)** Scheme depicting the anatomical organization of the recorded tdTomato+ SST neurons in CA1. **(B)** Micrograph showing a tdTomato+ SST neuron during whole-cell patch-clamp recording in CA1. Bar, 30 µm. **(C)** Examples of voltage changes and firing pattern of tdTomato+ SST neurons in Miro1^SST-WT^ and Miro1^SST-cKO^ mice responding to similar hyper– and depolarizing injection currents. **(D)** Averaged action potential frequency in control and Miro1^SST-cKO^ SST neurons following sequential depolarizing current injections (n= 7-9 cells per group obtained from at least 3 mice each; two-way ANOVA followed by Sidak’s test). **(E)** Examples of traces and spontaneous postsynaptic currents (sPSCs) in tdTomato+ SST neurons of Miro1^SST-WT^ and Miro1^SST-cKO^ mice clamped at –70 mV. **(F)** Quantification of sPSC frequency and amplitude in tdTomato+ SST neurons (n= 7-9 cells per group obtained from at least 3 mice each; unpaired t-test). **(G)** Scheme depicting the experimental setup used to record from pyramidal principal cells in CA1 in presence/absence of optogenetic stimulation of tdTomato/ChR2-expressing SST neurons. **(H)** Micrograph showing a recorded CA1 pyramidal neuron filled with biocytin, exemplifying the location of its apical dendrites with respect to axon terminals of nearby tdTomato+ SST neurons. Bar, 50 µm. **(I)** Examples of excitatory and inhibitory sPSCs recorded at –60 and 0 mV, respectively, in pyramidal CA1 neurons of Miro1^SST-WT^ and Miro1^SST-cKO^ mice. **(J)** Quantification of the E/I balance in recorded pyramidal CA1 neurons (n= 14-18 cells per group obtained from at least 3 mice each; unpaired t-test with Welch’s correction). **(K)** Quantification of the coefficient variability for pulses 2 to 5 during a 1 Hz optogenetic stimulation (1 ms) of SST+ neurons (n= 13-16 cells per group obtained from at least 3 mice each; unpaired t-test with Welch’s correction). **(L)** Examples and **(M)** quantification of c-Fos immunostaining in the CA1 and DG regions of Miro1^SST-WT^ and Miro1^SST-cKO^ mice (n= 3 mice per groups; Mann-Whitney test). **(N)** Heatmap of total amine-containing polar metabolites in the indicated genotypes (n= 6 mice per groups). Right bar reports on the significance of detected metabolites (– Log10 *P*-value ≥ 1.3). **(O)** Normalized fold change of GABA levels from dataset shown in N (n= 6 mice per groups; unpaired t-test). **(P)** Quantification of GABA and Vgat levels at axon terminals of tdTomato+ SST neurons (n= 3 mice per groups; Mann-Whitney test). *, p < 0.05, **, p < 0.01. S.o., stratum oriens; s.p., stratum pyramidale; s.r., stratum radiatum; s.l.m., stratum lacunosum-moleculare. See also Figure S5.

In net contrast, recordings from non-recombined, nearby CA1 pyramidal cells, which are innervated by axon terminals of SST+ neurons at their distal apical dendrites (Figures 5G-5H), revealed an overall reduced frequency of inhibitory sPSCs, leading to a significantly increased excitation/inhibition (E/I) input balance in Miro1^SST-cKO^ mice (Figures 5J). This suggests that while ablation of *Miro1* does not cell-autonomously overtly impact the membrane properties and synaptic inputs of recombined SST+ neurons, it does affect their inhibitory output onto principal cells. To verify this point, and assess reliability of synaptic transmission onto pyramidal cells, we virally transduced SST+ neurons with a Cre-dependent AAV encoding for ChR2-EYFP, and performed recordings of pyramidal cells in the transduced area during pulsed optogenetic field stimulation at 1 Hz (Figure 5G). This low-frequency stimulation protocol elicited stereotypical responses in pyramidal cells of Miro1^SST-WT^ mice, characterized by a progressive reduction in elicited peak amplitudes over pulses, yet the effect was visibly less pronounced in neurons of Miro1^SST-cKO^ mice (Figures S5J), which exhibited a higher variability coefficient between pulses (Figures 5K), suggesting a less reliable inhibition by SST+ neurons. Consistent with this finding, c-Fos immunoreactivity across CA1 and DG hippocampal fields was significantly elevated in 7-8-week-old Miro1^SST-cKO^ mice (Figures 5L-5M), denoting increased basal network excitability. Whole-hippocampus metabolomics at this age further revealed that besides a potential dysregulation of polyamine biosynthesis (spermidine and S-adenosyl methionine or SAM) and cysteine metabolism (cysteine, S-adenosyl methionine, methylthioadenosine or MTA, reduced glutathione or GSH, and homocysteine or Hcy), there was a partial but significant reduction in the concentration levels of the neurotransmitter GABA (minus 14% compared to control mice) right at the onset of seizure development, but not of many other amino acids including glutamate (from which GABA is primarily synthetized) (Figures 5N-5O). We used confocal imaging to spatially validate these findings, and found a meaningful decrease in GABA immunoreactivity within the axonal boutons of SST+ neurons in the *lacunosum-moleculare* at 8 weeks, while expression of the vesicular GABA transporter Vgat appeared only moderately affected (Figures 5P and S5K).

Collectively, these results provide evidence for a role of MIRO1 and mitochondrial transport in sustaining reliable SST+ neuron GABAergic transmission to balance hippocampal network excitability.

### Accumulation of dysfunctional synaptic mitochondria at the onset of seizures

While ablation of either *Miro1* or *Trak1* in SST+ neurons was sufficient to cause an hyperexcitable neuronal network, light microscopy proved ineffective in revealing any major alterations of mitochondrial structure at axon terminals (Figures 3C and 3G). We therefore used transmission electron microscopy (TEM) to investigate mitochondrial ultrastructure *in vivo*. To identify SST+ neurons in the CA1 *stratum oriens* of Miro1^SST-cKO^ mice, we took advantage of a CRE-dependent AAV expressing a genetically-encoded endoplasmic reticulum (ER)-targeted HRP (Joesch et al., 2016). Subsequent diaminobenzidine (DAB) staining enabled the identification of sparse SST+ neuronal cell bodies and dendrites among the much more numerous neurites of pyramidal cells (Figure 6A). We found that by 6 weeks of age, deletion of *Miro1* was sufficient to disrupt mitochondrial ultrastructure across HRP labelled soma and dendrites, with mitochondria appearing not only visibly rounder and enlarged in dendrites (corroborating the clusters visualized by light microscopy), but also containing regions with disorganized cristae, or altogether depleted of their characteristic cristae complexity otherwise seen in control neurons (Figure 6B). A noticeable formation of mitochondrial clusters and cristae disruption also occurred in CA1 and DG following CRE-dependent overexpression of SNPH in SST^tdTomato^ mice (Figure S6A and S6B), a paradigm which impairs mitochondrial motility (Kang et al., 2008), indicating that strategies blocking transport independently from MIRO1 manipulation are sufficient to cause ultrastructural mitochondrial defects in GABAergic SST+ neurons. Different from Miro1^SST-cKO^ mice, mitochondria in Miro1^PV-cKO^ mice exhibited a milder and much delayed ultrastructural phenotype which started becoming visible only around 14-16 weeks of age (Figure S6C), presumably reflecting a slower activation of the *Cre* transgene in the PV^Cre^ line or an altogether reduced basal turnover rate of PV+ compared to SST+ neurons (Figure 2F). Notably, the labelled ER in the neuronal soma of Miro1^SST-cKO^ and Miro1^PV-cKO^ mice appeared to retain close contact with wide portions of the mitochondrial outer membrane as in control neurons (Figures 6B and S6C), arguing against a primary role of MIRO1 in maintaining ER-mitochondria contact sites (Modi et al., 2019) in SST+ and PV+ neurons.

**Figure 6.**
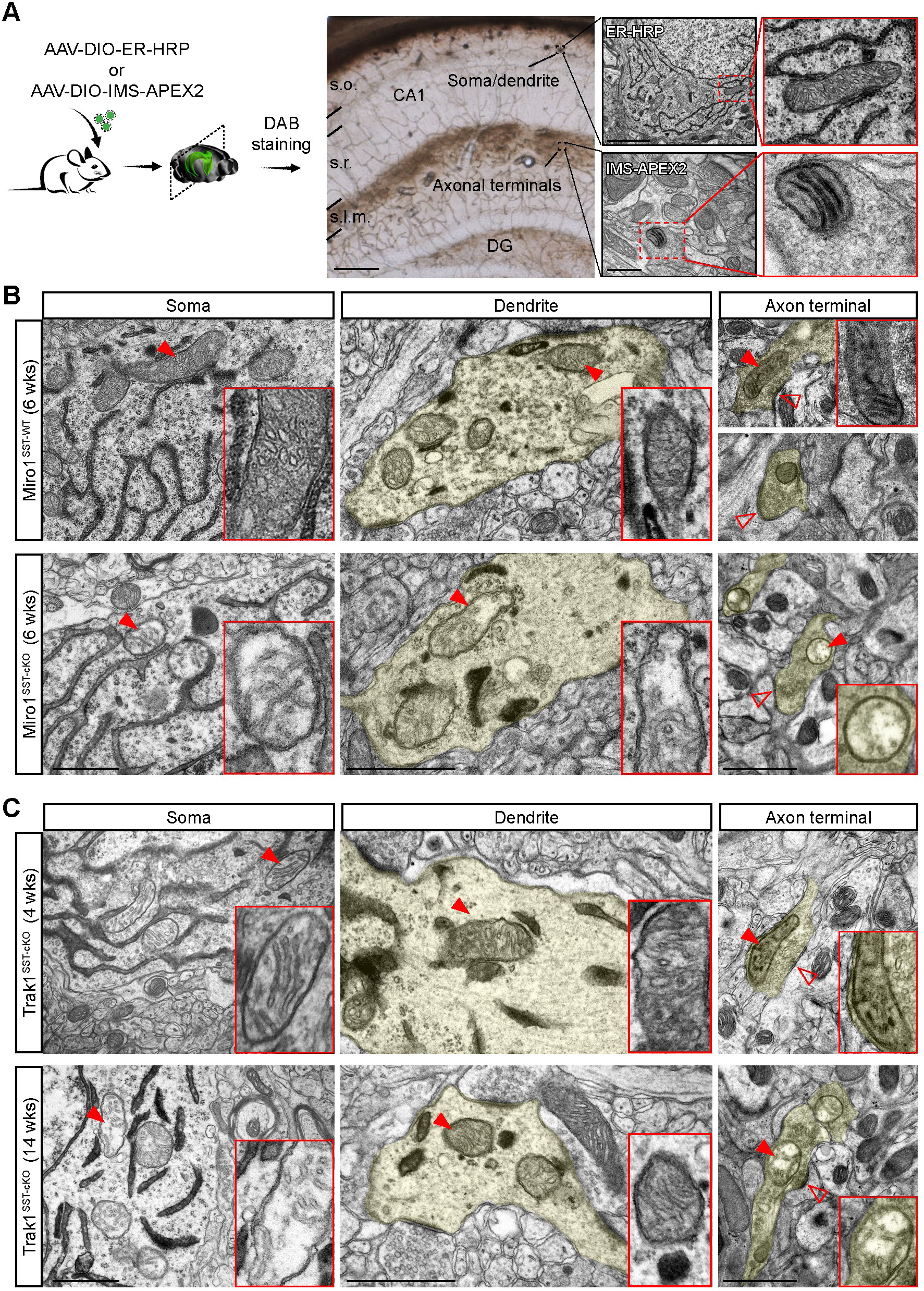
Accumulation of structurally dysfunctional mitochondria at axon terminals of SST+ neurons lacking *Miro1* or *Trak1*. (**A**) Experimental setup used for the Cre-dependent expression of ER-HRP or IMS-APEX2 in SST neurons, followed by DAB staining and EM. Right panels depict a hippocampal slice after DAB reaction, and examples of identified mitochondria within labelled cell bodies and axon terminals in the respective layers of CA1. Bars, 2 µm and 500 nm. **(B)** Electron micrographs of soma, dendritic region and axon terminals of labelled SST neurons in Miro1^SST-WT^ and Miro1^SST-cKO^ mice at 6 weeks of age. Insets show zooms of selected mitochondria (red arrowheads). Empty arrowheads point to putative synaptic contacts. Bars, 1 µm. **(C)** Electron micrographs of soma, dendritic region and axon terminals of labelled SST neurons in Trak1^SST-cKO^ mice at 4 and 14 weeks of age. Insets show zooms of selected mitochondria (red arrowheads). Empty arrowheads point to putative synaptic contacts. Bars, 1 µm. S.o., stratum oriens; s.r., stratum radiatum; s.l.m., stratum lacunosum-moleculare. See also Figure S6.

Given the apparent lack of significant ER-HRP labelling within axonal boutons of recombined SST+ neurons, we used conditional expression of a mitochondrial intermembrane space (IMS)-targeted APEX2 (Zhang et al., 2019) to specifically visualize mitochondria within axonal varicosities projecting to the *lacunosum-moleculare* (Figure 6A). There, we found numerous examples of labelled mitochondria in ultrastructurally intact putative synaptic boutons contacting unlabelled dendritic shafts, or their protrusions, in Miro1^SST-cKO^ mice. These contacts entailed the presence (at the labelled presynaptic site) of small and clustered vesicles, often exhibiting a mixture of round and flatten morphologies, a synaptic cleft, and a post-synaptic membrane lacking a prominent post-synaptic density (Figure 6B), which would be consistent with symmetric synapses (Kunkel et al., 1988). Interestingly, most presynaptic boutons contained one or two small mitochondria in both Miro1^SST-cKO^ and control mice (Figure 6B), corroborating our earlier observation that mitochondria were not depleted from the terminal by this stage (Figures 3E-3F). Consistently, there was no visible increase in autophagic/mitophagic vacuoles at these terminal sites (Figure 6B) or anywhere across the examined neuronal territories, as also evidenced by qualitative assessment of mitochondrial localization with lysosomal markers such as LAMP1 and LAMP2 (Figures S7A-S7B). In contrast to labelled mitochondria of Miro1^SST-WT^ mice, or within control unlabelled synapses in Miro1^SST-cKO^ mice, which alternated round with short tubular morphologies, mitochondria in MIRO1-deficient terminals exclusively exhibited a rounder morphology, and were almost completely devoid of cristae integrity (Figure 6B). Notably, the loss of mitochondrial cristae at axon terminals was more conspicuous than that of somato-dendritic mitochondria of the very same samples, where some remnants of disorganized cristae could be discerned (Figure 6B), supporting the notion that distal presynaptic sites experienced more profound consequences of disrupted mitochondrial turnover (Figure 2G). A remarkably similar ultrastructural phenotype was found also in axon terminals of Trak1^SST-cKO^ animals right at the onset of seizures (14 weeks), with mitochondria appearing with round morphologies and disrupted cristae, situated in otherwise preserved synaptic contacts (Figure 6C). Intriguingly, defects in mitochondrial ultrastructure in Trak1^SST-cKO^ mice appeared largely confined to the soma and axon, since mitochondria within dendrites presented with a much milder or absence of phenotype (Figure 6C), suggesting that these regionalized structural defects resulted from cumulative secondary effects caused by disruption of transport preferentially along the axon.

Together, these data show that the onset of seizures in Miro1^SST-cKO^ and Trak1^SST-cKO^ mice coincides with the occurrence of ultrastructural deficits in mitochondrial cristae of SST+ neurons, most prominent at their distal synaptic terminals, reinforcing the notion that the resulting neuronal network hyperexcitability originates from turnover defects at axonal presynaptic sites in these mice.

### Targeted gene therapy via systemic AAVs ameliorates the epileptic phenotype of Miro1^SST-cKO^ and Trak1^SST-cKO^ mice

Given the refractory nature of recurring seizures in patients carrying *TRAK1* pathogenic variants, which often leads to fatal outcomes (Barel et al., 2017; Li et al., 2024a; Sagie et al., 2018), we next sought to generate AAV vectors for preclinical therapeutic treatment of Miro1^SST-cKO^ and Trak1^SST-cKO^ mice. We first tested whether post-weaning *Miro1* re-expression via intracranial AAV delivery to CA1 would ameliorate morpho-functional aspects of mitochondria in Miro1^SST-cKO^ mice, examining morphology and membrane potential (Figure 7A). A period of 2 weeks after injecting an AAV engineered to express *Miro1* in combination with (or without) *mtGFP* (AAV-flex-Miro1-T2A-mtGFP, referred to AAV-Miro1) into 4-week-old animals was sufficient to largely restore a tubular network in the soma and dendrites of MIRO1-deficient neurons (Figure S7C). Likewise, membrane potential as evaluated by 2PLSM and Tetramethylrhodamine methyl ester (TMRM) staining of acute brain slices revealed a marked reduction in SST+ neurons of Miro1^SST-cKO^ compared to control mice, while *Miro1* re-expression led to a significant recovery (Figure 7B-7C). This indicates that morphological and functional mitochondrial parameters can be rapidly reversed by *Miro1* re-expression *in vivo* in young animals.

**Figure 7.**
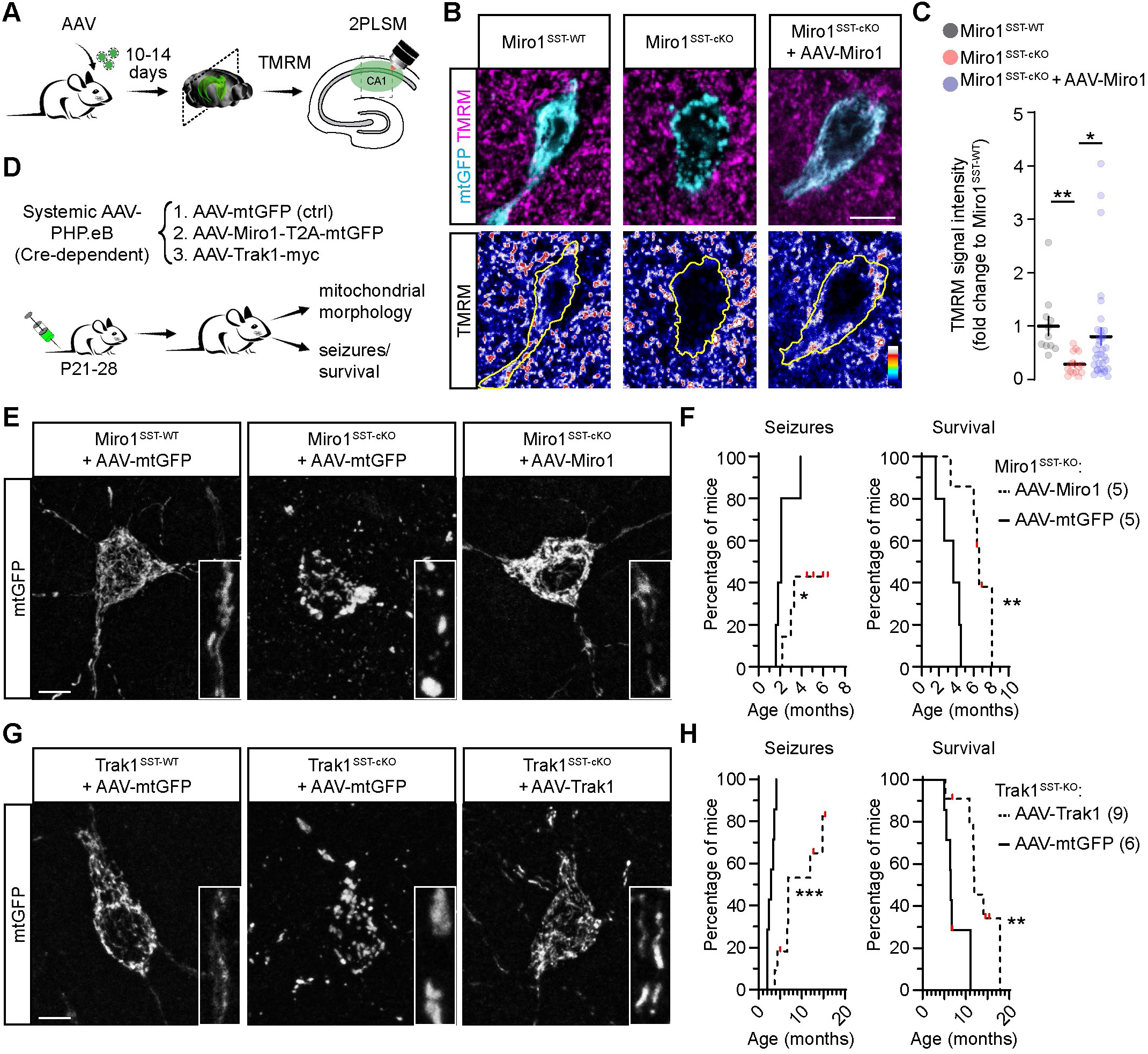
Targeted post-natal re-expression of *Miro1* and *Trak1* ameliorates the epileptic phenotype and improves survival of Miro1^SST-cKO^ and Trak1^SST-cKO^ mice. (**A**) Experimental setup used for the Cre-dependent expression of Miro1-T2A-mtGFP in SST neurons of Miro1^SST-WT^ and Miro1^SST-cKO^ mice, followed by TRMR staining and 2PLSM imaging in acute hippocampal slices. **(B)** Examples of mtGFP-expressing SST neurons in slices stained with TMRM. Lower panels show pseudocolored TMRM signal intensity levels. Bar, 10 µm. **(C)** Quantification of TMRM signal intensity for the conditions shown in B (n= 11-34 cells, obtained from at least 3 mice per condition; Kruskal-Wallis followed by Dunn’s test). **(D)** Experimental setup used for the Cre-dependent expression of systemic PHP.eB AAVs expressing mtGFP alone, Miro1-T2A-mtGFP or Trak1-myc in mice after weaning. **(E)** Examples of mtGFP-expressing hippocampal CA1 SST neurons in Miro1^SST-WT^ and Miro1^SST-cKO^ mice after systemic AAV injection. Insets report on zooms of dendritic regions. Bar, 10 µm. **(F)** Graphs depicting the incidence of seizures and survival rate over time Miro1^SST-cKO^ mice after systemic infusion of control AAV or AAV-Miro1 (n= 5 mice per group; Mantel-Cox test). **(G)** Examples of mtGFP-expressing hippocampal CA1 SST neurons in Trak1^SST-WT^ and Trak1^SST-cKO^ mice after systemic AAV injection. Insets report on zooms of dendritic regions. Bar, 10 µm. **(H)** Graphs depicting the incidence of seizures and survival rate over time Trak11^SST-cKO^ mice after systemic infusion of control AAV or AAV-Trak1 (n= 6-9 mice per group; Mantel-Cox test). *, p < 0.05, **, p < 0.01, ***, p < 0.001. See also Figure S7.

We next assessed the efficiency of SST+ cell recombination upon administration of a CRE-dependent (flexed), systemic AAV bearing a capsid variant (PHP.eB) suitable for crossing the mouse blood-brain barrier (Chan et al., 2017) (Figure 7D). We specifically focused on delivering and testing the effects of systemic AAVs at post-weaning stages, reasoning that this would realistically model a time window suitable for therapeutic applicability. A single tail vein injection was sufficient to elicit a transduction rate of 53% and 49% in the neocortex and hippocampus, respectively, as assessed by AAV reporter (mtGFP) expression over the endogenous tdTomato reporter encoded by the SST^tdTomato^ line (Figure S7D-S7E). Like for the intracranially delivered AAV, *Miro1*-expression via systemic AAV injection was sufficient to significantly improve mitochondrial morphology across the somato-dendritic compartment in Miro1^SST-cKO^ mice, in contrast to neurons transduced with the control AAV, which retained a fragmented and clustered mitochondrial network (Figure 7E). This effect was accompanied by a prominent delay in the occurrence of seizures and a corresponding significant extension in survival, with a median of 6.6 months in mice treated with AAV-Miro1 versus 3.6 months in the control group (Figure 7F). Similarly, a single systemic administration of a *Trak1*-expressing AAV was sufficient to re-establish a more physiological mitochondrial morphology in transduced SST+ neurons (Figure 7G), significantly delaying the onset of seizures and almost doubling the median survival of Trak1^SST-cKO^ mice from 6.5 months (control group) to 11.9 months (treated group) (Figure 7H).

Thus, systemic AAV administration for targeted *Miro1* and *Trak1* expression has preclinical applicability in ameliorating the epileptic phenotype and improving the survival rate of mutant mice.

## Discussion

We provide here evidence for experience-dependent changes in mitochondrial proteome and relative turnover rates across major hippocampal neuronal subtypes, identifying a distinctive response in GABAergic neurons – particularly SST+ ones – at their distal presynaptic sites, underscoring a cell type-specific role for axonal mitochondrial transport in maintaining neuronal network excitability.

Mitochondrial turnover is widely considered key for neuronal integrity, yet dissecting lifetimes of single organelles, their protein or lipid content across neuronal types and sub-compartments *in vivo* has proven challenging. For instance, the use of photoswitchable optical sensors has only recently enabled to infer a mitochondrial lifetime of about 3-5 days in large motor neuron synaptic terminals, demonstrating that this relatively short turnover rate depends on coordinated axonal transport dynamics and local degradation mechanisms (Marahori et al., 2024). Such timeline would be consistent with the rapid experience-dependent changes in mitoTimer signal that we detected in the comparatively much smaller hippocampal cell types examined here, especially considering that our analysis exclusively focused on relative turnover rates, as mitoTimer precludes a direct assessment of absolute mitochondrial lifetime (Trudeau et al., 2014). While turnover rates of days may be compatible with previously quantified half-life of mitochondrial proteins in cultured neurons (Dorrbaum et al., 2018), *in vivo* metabolic isotope labelling revealed the existence of exceptionally long-lived (from weeks up to several months) subsets of mitochondrial proteins in brain tissue (Bomba-Warczak et al., 2021; Fornasiero et al., 2018), emphasizing the existence of multi-layered quality control mechanisms that contribute to maintain mitochondrial network homeostasis, possibly in a cell type-specific manner, across the lifespan. Understanding how these dynamics may be regulated *in vivo* is further complicated by the fact that experience and circuit activity states are likely to modify mitochondrial protein composition (Alvarez-Castelao et al., 2017; Fornasiero et al., 2018; Heo et al., 2018; McNair et al., 2007), as these depend on a multitude of intertwined processes (biogenesis, degradation, transport, fission-fusion dynamics) whose relative contribution also compartmentalizes across extended neuronal geometries to serve local metabolic needs (Cawley and Farris, 2026; Devine and Kittler, 2018). Supporting this notion, there is mounting evidence for molecular and functional specialization of mitochondria across sub-cellular compartments, cell types and brain regions, implying that this phenotypic diversification plays important roles in circuit activity and higher brain functions (Fecher et al., 2019; Hirabayashi et al., 2024; Mosharov et al., 2025; Pastor et al., 2025; Rosenberg et al., 2023).

By comparatively examining the soma and axon terminals of various neuronal classes in the mouse hippocampus, we identified important cell type-specific differences between glutamatergic and GABAergic neurons induced by a novel EE. Within the cell body, where the bulk of mitochondrial biogenesis and degradation are thought to occur, the mito-proteome of glutamatergic neurons was left largely unchanged after EE, while the relative abundance of many proteins (including several OxPhos subunits) in GABAergic neurons was significantly decreased, leading to an overall specific shift of the mitochondrial versus whole-cell proteomes (Figure S1F). While our proteomics approach prevented us to reach any conclusions about changes in protein half-lives, or their relative abundance in dendritic versus axonal compartments (which were both excluded from FACS), it retained cell type specificity, and painted a picture consistent with an experience-dependent downregulation of several pathways associated to energy metabolism in PV+ and SST+ neurons. The underlying mechanism of this selective response of the mitochondrial proteome in PV+ and SST+ neurons remains unclear. Given that the activity of both classes of GABAergic neurons in CA1 is rapidly suppressed during spatial exploration of novel environments (Arriaga and Han, 2019; Geiller et al., 2020; Hainmueller et al., 2024) to lessen network inhibition and promote plasticity of excitatory place cells (Sheffield et al., 2017; Udakis et al., 2025), it is tempting to speculate that the changes in mito-proteome seen here within the soma of PV+ and SST+ neurons may reflect a switch of activity state. This may then be enabled via activity-dependent changes at transcriptional and translational levels (Li et al., 2024b), but possibly also by recruiting additional quality control mechanisms that expedite mito-proteome rewiring to rapidly cope with the new metabolic demands, as shown for other types of neural cells (Wani et al., 2022).

Synchronized, pulse-chase mitoTimer experiments in the same EE settings showed a substantially higher proportion of redder mitochondria within the soma compartment of glutamatergic versus GABAergic neurons, while the fraction of highly green mitochondria (i.e., reflecting increased biogenesis) was unchanged across cell types. Given the spontaneous mitoTimer maturation rate from green to red by 48h after the first Dox pulse (Trudeau et al., 2014), which was extended here to a 3-day-long period for normalization purposes across examined cell types, the specific accumulation of mitochondria in the redder fraction by 1 day after the last Dox pulse likely reflects a reduced degradation rate of “older” mitoTimer in the soma of glutamatergic neurons, similar to what has been described following autophagy ablation experiments (Ferree et al., 2013). While we can only speculate on the potential differences in activity-dependent degradative mitochondrial mechanisms between the examined cell types, our mitoTimer data would support a comparatively higher mitochondrial turnover rate in the soma of GABAergic than glutamatergic neurons after EE.

In contrast to mitochondria located in the perisomatic region, their positioning at distal axon terminals is regulated by long-range transport dynamics (Misgeld and Schwarz, 2017), which together with local degradation and translation mechanisms account for the bulk of mitochondrial turnover at distal axons (Cioni et al., 2019; Maday et al., 2012; Marahori et al., 2024). We indeed found disjoint mitoTimer dynamics between perisomatic and axon terminal regions in the very same cells types, with a distinctive effect in SST+ neurons towards a reduced relative turnover rate induced by novel EE. While our analysis revealed that this effect was due to a remarkably higher proportion of greener mitochondria at the terminals of SST+ neurons compared to other neuronal types, reflecting an enhanced turnover rate at basal conditions, how this pronounced sensitivity to a novel EE is coordinated at synaptic sites of SST+ neurons remains unclear. Importantly, this effect was independent from the absolute distance between synaptic terminals and the neuronal cell body, as a role played by shorter distances would be mirrored by even higher turnover rates in PV+ GABAergic neurons (e.g., basket cells), which was not the case. Supporting a specific role for axonal mitochondrial transport in this process, experimental blockade of transport dynamics in Miro1^SST-cKO^ mice was sufficient to shift synaptic MitoTimer signal towards a redder range, with accumulating mitochondria undergoing loss of cristae at a much higher pace than in Miro1^PV-cKO^ mice, indicating sustained mitochondrial motility in mature SST+ neurons. Interestingly, the same analysis revealed opposite basal turnover dynamics at the axon terminals of glutamatergic CA1 pyramidal and DG granule neurons, as both cell types exhibited a large fraction of older mitochondria which appeared largely insensitive to EE (Figure 2F), consistent with the previously reported very low rates of mitochondrial transport in mature upper layer callosal projection axons (Lewis et al., 2016; Smit-Rigter et al., 2016). Hence, while we cannot rule out that specialized sites of mitochondrial biogenesis or degradation may also play a role in mitoTimer dynamics at SST+ neuron terminals, their enhanced mitochondrial motility rates may represent a neuron subtype-specific trait dictated by anatomical and functional constrains, including differences in basal firing activity, axon calibre and myelination degree (Call and Bergles, 2021; Stedehouder et al., 2019), which together play important roles in defining mitochondrial distribution rules at synaptic terminals (Kole et al., 2022; Macaskill et al., 2009; Ohno et al., 2011; Silva et al., 2021; Wang and Schwarz, 2009), and thus likely also contribute to define mitochondrial phenotypic diversity between neuronal cell types (Sager et al., 2026). Extending compartmentalized analyses of mitochondrial turnover dynamics to other disease-relevant neuronal populations, such as dopaminergic neurons, may therefore provide insights into how defective mitochondrial transport and/or function would confer selective synaptic vulnerabilities, as exemplified by the growing evidence for a role of MIRO1 and its mutations in Parkison’s disease (Chemla et al., 2025; Hsieh et al., 2019; Wang et al., 2011).

Strikingly, all GABAergic neuron-specific mouse lines used in our study to ablate MIRO1 and TRAK1 did not show visible signs of synaptic or neuronal degeneration at any of the examined time points, in contrast to models of forebrain *Miro1* deletion (López-Doménech et al., 2016; Nguyen et al., 2014), and despite the emergence of recurrent seizures in Miro1^SST-cKO^ and Trak1^SST-cKO^ mice, as well as hypertonia linked to peri/postnatal lethality in Miro1^Vgat-cKO^ and Trak1^Vgat-cKO^ mice. These phenotypic traits are highly reminiscent of those reported in the hypertonic (*hyrt*) mouse model found to be constitutively mutated in the mouse *Trak1* gene (Gilbert et al., 2006), and in patients carrying *TRAK1* loss-of-function pathogenic variants, which manifest as congenital hyperekplexia/myoclonus, and early-onset seizures progressing to refractory status epilepticus during infancy, followed by death (Barel et al., 2017; Li et al., 2024a; Sagie et al., 2018). Fibroblasts derived from these patients exhibit disrupted mitochondrial distribution, motility, and membrane potential (Barel et al., 2017), which are features also recapitulated in our mouse models. However, SST neuron axon terminals of Miro1^SST-cKO^ and Trak1^SST-cKO^ mice were not depleted of mitochondria, different from previous studies conducted in other species (Guo et al., 2005; Stowers et al., 2002) or in mice undergoing early-onset recombination (López-Doménech et al., 2016; Nguyen et al., 2014). This discrepancy may thus reflect either a GABAergic cell type-specific phenotype or a developmentally delayed recombination efficiency of the SST^Cre^ driver line used here compared to constitutive mutants. This delay would result in complete MIRO1 (or TRAK1) protein ablation only at relatively late stage of postnatal maturation, when GABAergic neuron migration and arborization, alongside initial mitochondrial positioning, may have been already initially achieved (Lim et al., 2018), and then needs to be maintained. Alternatively, mitochondrial depletion in our models may become evident only at times later than those examined here, when mitochondrial dysfunction would become even more severe. However, assessing mitochondrial occupancy within distal terminals at longer time points would be ethically challenging to perform given the premature lethality of the used mouse models. Yet, the preserved integrity of synapses and neurons in Miro1^SST-cKO^ and Trak1^SST-cKO^ mice up to seizure onset suggests a temporal sequence where mitochondrial dysfunction emerges as consequence to a faulty transport, but is primarily responsible for the metabolic deficit driving reduced GABAergic transmission. This sequence is supported by the following facts: (i) conspicuous deficits in axonal mitochondrial transport were the earliest to occur at 3 weeks of age as seen by live imaging experiments, while disrupted mitochondrial morphology took place starting from 4 weeks and later times points; (ii) mitochondrial ultrastructural deficits were most prominent at distal axon terminals, where turnover is anticipated to be most affected by blockade of transport compared to cell body; and (iii), seizures in Miro1^SST-cKO^ and Trak1^SST-cKO^ mice only kicked-in upon significant cristae disruption, emphasizing the underlying role of mitochondrial dysfunction in this phenotype. In these settings, the reduced GABAergic transmission may therefore result from local depletion of key metabolites (i.e., ATP supply) or dysregulation of Ca2+ buffering, which are processes required for maintenance and activity-dependent release of synaptic vesicle pools (Ashrafi et al., 2020; Pulido and Ryan, 2021; Rangaraju et al., 2014; Verstreken et al., 2005). Alternatively, defective neurotransmission may be caused by a decreased bioavailability of synaptic GABA itself, which depends on mitochondria-derived glutamate for its own biosynthesis on one hand, and on the other hand on mitochondrial import to feed the TCA cycle and sustain respiration (so-called GABA shunt) (Balazs et al., 1970; Hassel et al., 1998), a process that consumes GABA and leads to neurological disorders when dysregulated (Jaeken et al., 1984; Kanellopoulos et al., 2020; Pearl et al., 2011). Consistent at least in part with the latter possibility, Miro1^SST-cKO^ mice exhibited a partial but meaningful reduction in their hippocampal GABA levels (Figures 5N-5P), reflecting the genetic recombination restricted to SST+ neurons, which themselves account for about 15-20% of the entire hippocampal GABAergic neuron population (Freund and Buzsaki, 1996).

Mitochondrial dysfunction has been long recognized as one possible source of epilepsy (Zsurka and Kunz, 2015). In our study we showed that mice lacking *Miro1* in SST+ neurons display abnormal ECoGs, including interictal spikes and typical epileptic discharges. We assume that such abnormal network activity was caused by reduced GABA-mediated neurotransmission, which we demonstrated by patch-clamp recordings of principal neurons in CA1, and which is in line with the detrimental effect caused by null mutations in *Drosophila* dMiro for neurotransmitter release (Guo et al., 2005). Interestingly, power spectrum analysis revealed a significantly lower frequency in the theta band and amplitude in the gamma bands in mutant mice compared to controls. On one hand, these results may appear paradoxical, as an increase in the ECoG spectral amplitude could be expected when GABAergic neurotransmission is compromised. However, since inhibitory neurons are involved in the generation of oscillations by synchronizing neural network activity (Cardin, 2018), we speculate that the decreased power in mutant mice results from reduced synchronism, which can be caused by dysregulated inhibitory network function. On the other hand, our finding of a reduced frequency in theta band activity broadly supports previous work linking SST+ neuron activity with theta oscillation (Huang et al., 2020; Varga et al., 2012). One unanticipated finding was that gamma band oscillations were affected as well, raising the question whether *Miro1*-deficient SST+ neurons impact high-frequency oscillations directly acting on principal cells (Veit et al., 2017) or indirectly by acting on PV+ neurons, which are well-known to play a pivotal role in the generation of gamma oscillations (Hu et al., 2014).

Strategies to enhance mitochondrial transport have been adopted to improve the overall homeostasis of mitochondria, particularly following injury and in disease models. For example, boosting axonal mitochondrial trafficking via ablation of the docking protein SNPH or other means helps removing damaged mitochondria, improving axon regeneration (Cartoni et al., 2016; Han et al., 2020; Han et al., 2016; Lin et al., 2017). In our model, postnatally restoring mitochondrial movement by targeted *Miro1* or *Trak1* supplementation rescued mitochondrial integrity in transduced neurons, and considerably improved life expectancy of the epileptic mutant mice. These findings add to our understanding of how modulating mitochondrial transport may compensate for, or ameliorate axonal and synaptic metabolic deficits underlying disease, and have important implications for the potential use of systemic AAV delivery to restore gene expression in a cell type-specific manner.

## Author Contributions

Conceptualization, K.N. and M.B.; Methodology, K.N, A.C., M.J., P.P., V.S., F.G., A-L.S., E.M., S.E.T., M.A., D.I. and M.B.; Investigation, K.N., K.G., K.M.T., G.A.W., P.P., J.S.C., F.O., V.S., C.A.F., L.P.R., J.B., F.G., M.A., D.I. and M.B.; Formal analysis, K.N., A.C., K.G., K.M.T., G.A.W., P.P., V.S., A-L.S., F.O., L.P.R., S.M., M.A., D.I. and M.B.; Writing – Original Draft, M.B.; Writing – Review and Editing, K.N., A.C., K.G., M.J., K.M.T., G.A.W., P.P., J.S.C., F.O., V.S., C.A.F., L.P.R., J.B., A-L.S., F.G., A.S., E.M., S.E.T., B.Z., S.M., M.A., D.I. and M.B.; Resources, K.N, M.J., S.E.T., B.Z. and M.B.; Funding Acquisition, M.B.; Supervision, M.B.; Project Administration, M.J. and M.B.

## Supporting information

Key resource table

## Acknowledgements

We thank N.G. Larsson and J. Shaw for providing *mtYFP* flox-stop and *Miro1* floxed mice. C. Kukat, C. Jüngst, P. Zentis, V. Teiwes, P. Scharwächter and all the members of the core facilities in CECAD (in vivo Research Facility and Transgenic Core Unit, proteomics and imaging) and Max Planck Institute for Biology of Ageing (FACS & Imaging Core Facility) for excellent assistance. A. Kukat, M. Veronese, H.M. Jahn-Kelleter, G. Piper and all alumni of the Bergami lab for help on ethical and technical matters, general support and discussions. This work was supported by the Deutsche Forschungsgemeinschaft (SFB1218 – Grant No. 269925409, projects A07 and A11; and CECAD EXC 2030 – Grant No. 390661388 to M.B. and E.M; SFB1451 – Grant No. 431549029, projects A03 and B01 to M.B. and D.I.); the European Research Council (ERC-StG-2015, grant number 677844), and the Alzheimer Forschung Initiative e.V. (#24019R, with kind support of the Stiftung Alzheimer Initiative SAI) to M.B.; and by the Research Committee of the Medical Faculty, Heinrich Heine University (KS 9772904) to M.A.. L.P.R is supported by the Deutsche Forschungsgemeinschaft Walter-Benjamin program (Grant No. PE4191/1-1). K.M.T. is supported by the Jürgen Manchot Stiftung. Large instrumentation used in this work was supported by the Deutsche Forschungsgemeinschaft (DFG Großgeräteantrag): INST 1856/71-1 FUGG, INST 216/793-1 FUGG, INST 216/741-1 FUGB, INST 2016/742-1 FUGB.

## Legends to Supplementary figures

**Supplementary Figure 1.**
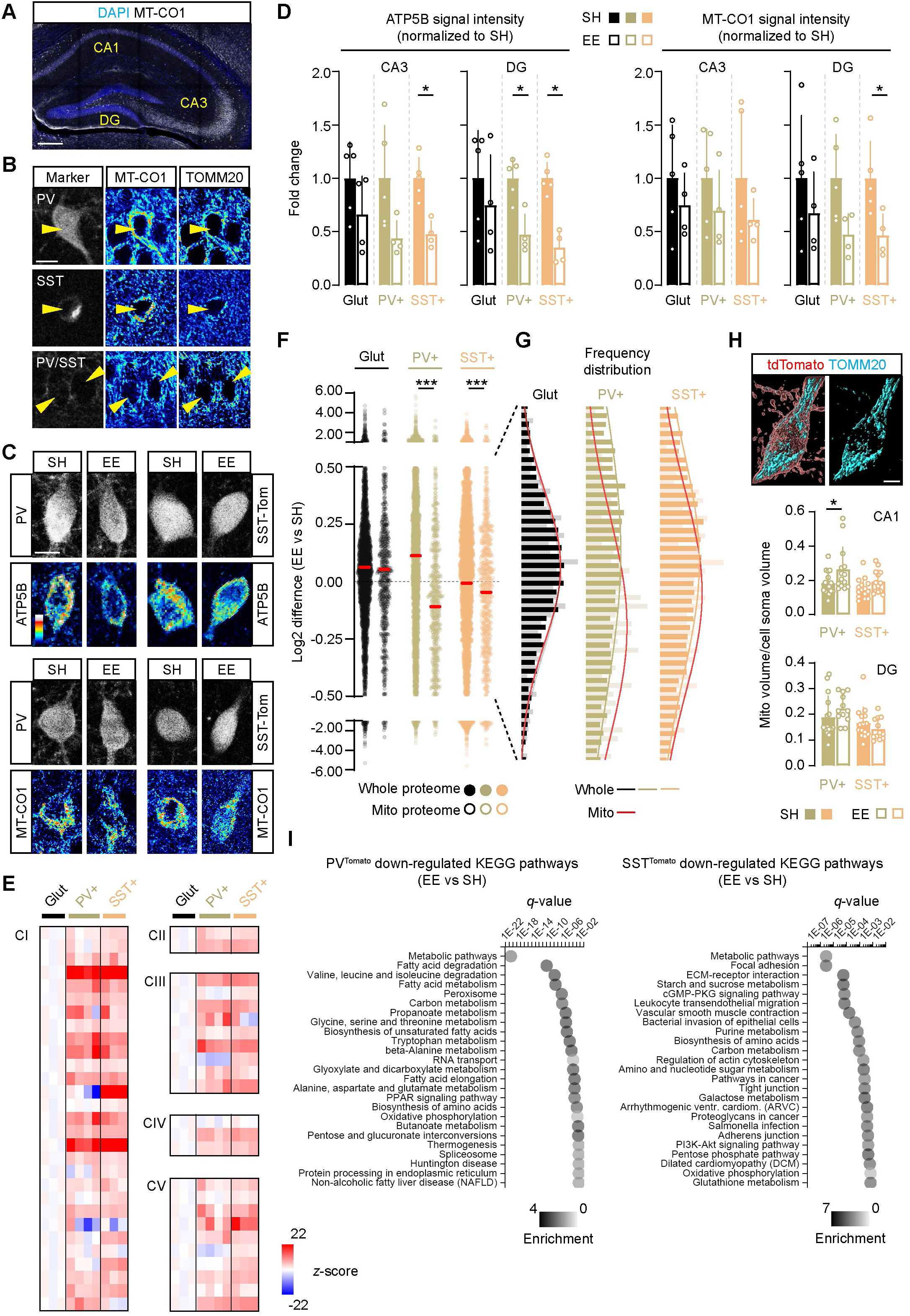
Selective experience-dependent rewiring of the mitochondrial proteome in GABAergic neurons. **(A)** Example of MT-CO1 immunostaining across hippocampal subfields. Bar, 80 µm. **(B)** Example of pseudocolored MT-CO1 and TOMM20 expression levels in PV+, SST+ and glutamatergic principal neurons in CA1. Bar, 8 µm. **(C)** Example of pseudocolored ATP5B (top panels) and MT-CO1 (lower panels) expression levels in CA1 PV+ and tdTomato+ neurons of SST^tdTomato^ mice maintained in SH or exposed to EE for 48h. Bar, 7 µm. **(D)** Quantification of the changes in ATP5B and MT-CO1 signal intensity for glutamatergic, PV+ and SST+ neurons in CA3 and DG regions after EE exposure (n= 4-5 mice per condition; unpaired Mann-Whitney test). **(E)** Heat-maps of relative abundance for respiratory complex subunits quantified in sorted glutamatergic, PV+ and SST+ neurons under SH conditions (n=3-4 mice per condition). **(F)** Scatter dot plot comparison of whole vs mito-proteome in glutamatergic, PV+ and SST+ neurons after EE exposure (n=3-4 mice per condition; Kolmogorov-Smirnov test). **(G)** Zoom of the central values (0.5 to –0.5 log2 difference) shown in F, reporting on the frequency distribution of whole (dark colors) vs mito-proteome (light colors and red connecting line) for the examined neuronal cell types. **(H)** Example of reconstructed PV+ neurons in CA1 showing the extent of mitochondrial occupancy (TOMM20 signal) over cell body volume. Bar, 5 µm. Lower graphs report on the quantification of mitochondrial mass over cell body volume for PV+ and SST+ neurons of CA1 and DG in SH and EE conditions (n= 12-15 cells over 4-5 mice per condition; unpaired two-tailed Mann-Whitney test). **(I)** Top 25 most significantly down-regulated KEGG pathways in PV+ and SST+ neurons after EE (Fisher exact test with FDR < 0.02, calculated on down-regulated proteins with *q* ≤ 0.05; n= 4-5 mice per condition). *, p < 0.05, ***, p < 0.001. S.o., stratum oriens; s.p., stratum pyramidale; s.r., stratum radiatum. See also Figure 1.

**Supplementary Figure 2.**
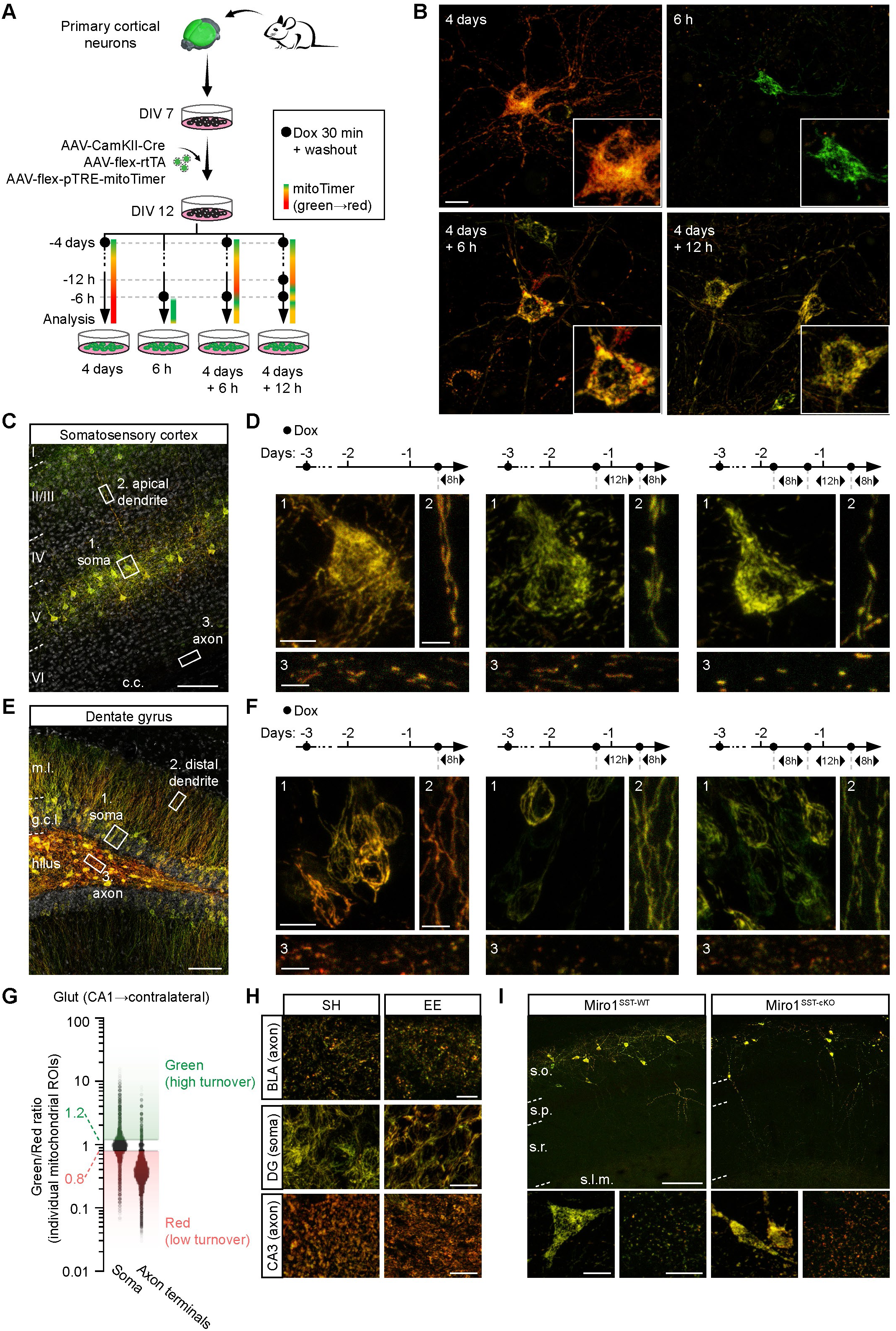
*In vitro* and *in vivo* validation of Dox-dependent mitoTimer dynamics. **(A)** Scheme showing the experimental setup used to test mitoTimer dynamics in neuronal cultures over 4 days. Five days after initial AAV transduction, a single 30 min-long pulse of Dox (black dots) was given to enable expression, followed by different protocols of chasing and/or additional Dox pulses. Four days later, neurons were analyzed. **(B)** Representative examples of each of the four conditions shown in A, illustrating the changes in green/red ratio of the mitochondrial network in transduced neurons at various times after one or multiple Dox pulses. Bar, 10 µm. **(C)** Example of mitoTimer-transduced neurons in the somatosensory cortex of wild-type mice. Bar, 70 µm. **(D)** Examples of cortical neurons as shown in C, depicting the changes in mitoTimer green/red ratio achieved in each cellular compartment by increasing the number of Dox pulses before analysis. Bars, 7, 5 and 3 µm. **(E)** Example of mitoTimer-transduced neurons in the DG of wild-type mice. Bar, 50 µm. **(F)** Examples of DG granule neurons as shown in E, depicting the changes in mitoTimer green/red ratio achieved in each cellular compartment by increasing the number of Dox pulses before analysis. Bars, 10, 5 and 3 µm. **(G)** Plots showing the green/red ratio distribution of all mitochondrial ROIs analyzed in a single field of view containing labelled glutamatergic neurons as shown in D. Cut-offs for high-green and high-red ratios are shown. Note the visibly redder signal (low turnover) in axon terminals as compared to their somatic counterpart. **(H)** Representative examples of mitoTimer green/red signal ratio in the indicated regions in SH or after EE. Bar, 12 µm. **(I)** Representative overviews and zooms of mitoTimer expression in CA1 of Miro1^SST-WT^ and Miro1^SST-cKO^ mice, showing the green/red merged signal for cell body and axon terminals of SST+ neurons. Bar, 70 µm. M.l., molecular layer; g.c.l., granule cell layer; See also Figure 2.

**Supplementary Figure 3.**
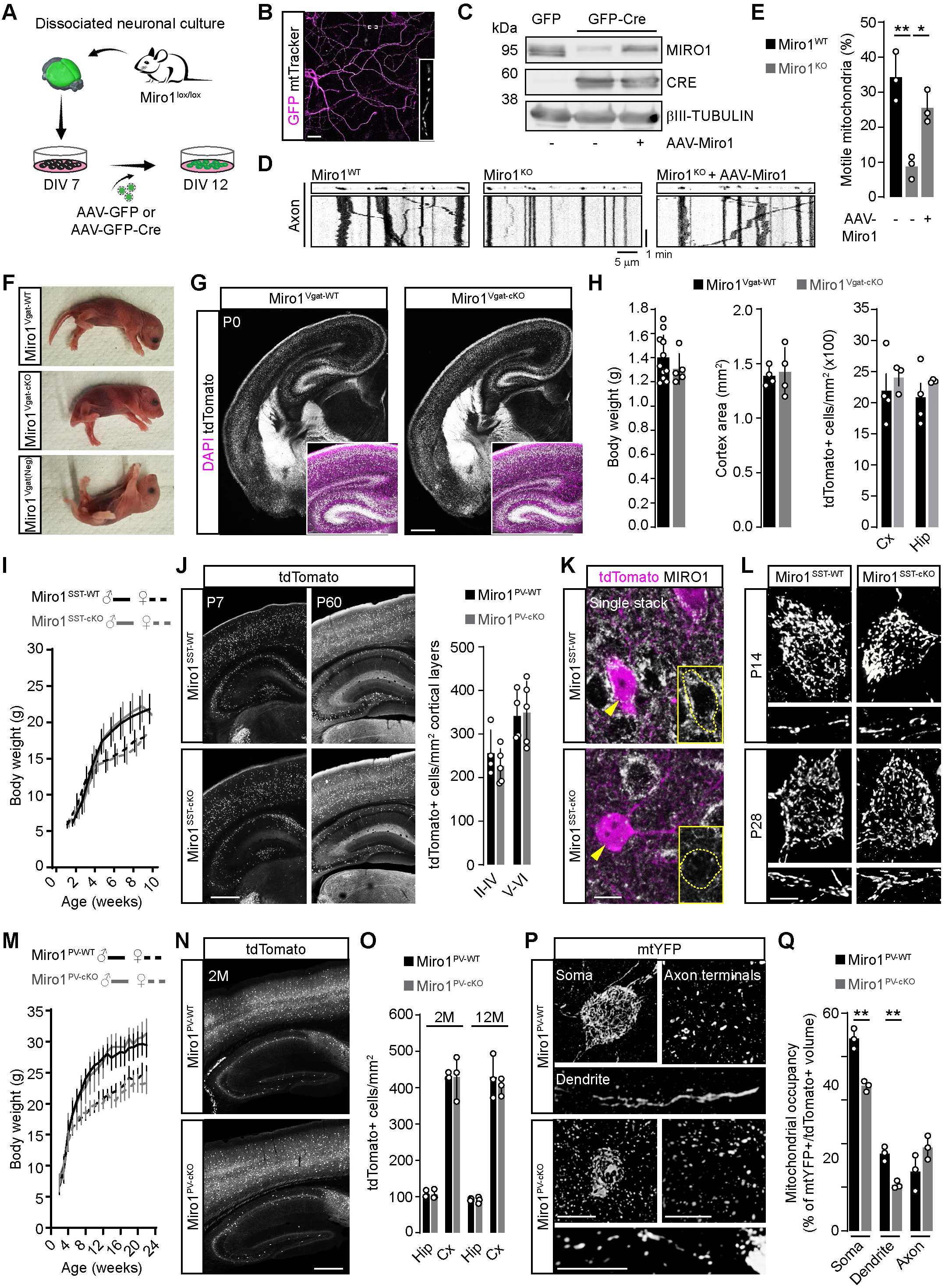
Phenotypic analysis of mice lacking *Miro1* in GABAergic neurons types. **(A)** Scheme showing the experimental timeline used to induce knockout of *Miro1* in neuronal cultures of Miro1floxed mice. **(B)** Example of AAV-transduced neurons in culture and **(C)** corresponding WB analysis of MIRO1 expression levels after application of GFP control or GFP-Cre-expressing AAVs, in presence or absence of AAV-Miro1. Bar, 15 µm. **(D)** Kymograph analysis of mitochondrial motility upon labelling with mitoTracker Red in axons in the same conditions as in C. **(E)** Quantification of axonal mitochondrial motility as shown in D (n= 3 experiments; one-way ANOVA followed by Tukey’s test). **(F)** Pictures showing Miro1^Vgat-KO^ and control pups alive at birth. **(G)** Examples of Miro1^Vgat-WT^ and Miro1^Vgat-KO^ brain sections at P0 expressing tdTomato. Bar, 500 µm. **(H)** Quantification of body weight (n= 8-11 pups per group; unpaired t-test), neocortex area (n= 4 pups per group; unpaired t-test) and tdTomato+ cell density (neocortex and hippocampus; n= 3-4 pups per group; two-way ANOVA followed by Sidak’s test) in Miro1^Vgat-WT^ and Miro1^Vgat-KO^ mice at P0. **(I)** Growth curve of Miro1^SST-WT^ and Miro1^SST-cKO^ mice up to 10 weeks of age (n= 3-16 mice per time point and group; mixed effect model followed by Bonferroni’s test). **(J)** Examples of Miro1^SST-WT^ and Miro1^SST-cKO^ brain sections expressing tdTomato at P7 and P60. Right plots depict density of tdTomato+ neurons across cortical layers at P60. Bar, 500 µm. **(K)** Immunostaining against MIRO1 in Miro1^SST-WT^ and Miro1^SST-cKO^ brain sections. Bar, 10 µm. **(L)** Examples illustrating the mitochondrial morphology within soma and dendrites of mtYFP-expressing neurons in the CA1 of Miro1^SST-WT^ and Miro1^SST-cKO^ mice at P14 and P28. Bar, 5 µm. **(M)** Growth curve of Miro1^PV-WT^ and Miro1^PV-cKO^ mice up to 24 weeks of age (n= 3-32 mice per time point and group; mixed effect model followed by Bonferroni’s test). **(N)** Examples of Miro1^PV-WT^ and Miro1^PV-cKO^ brain sections expressing tdTomato at P60. Bar, 400 µm. **(O)** Quantification of cortical and hippocampal tdTomato+ neurons at 2 and 12 months in Miro1^PV-WT^ and Miro1^PV-cKO^ mice (n= 3 mice per condition; two-way ANOVA followed by Sidak’s test). **(P)** Examples illustrating the mitochondrial morphology within soma, dendrites and axon terminals of mtYFP-expressing neurons in the CA1 of Miro1^PV-WT^ and Miro1^PV-cKO^ mice at P60. Bars, 8, 8 and 4 µm. **(Q)** Quantification of mitochondrial occupancy (over tdTomato+ cell volume) across neuronal compartments as shown in P (n= 3 mice per condition; unpaired t-test). *, p < 0.05, **, p < 0.01. See also Figure 3.

**Supplementary Figure 4.**
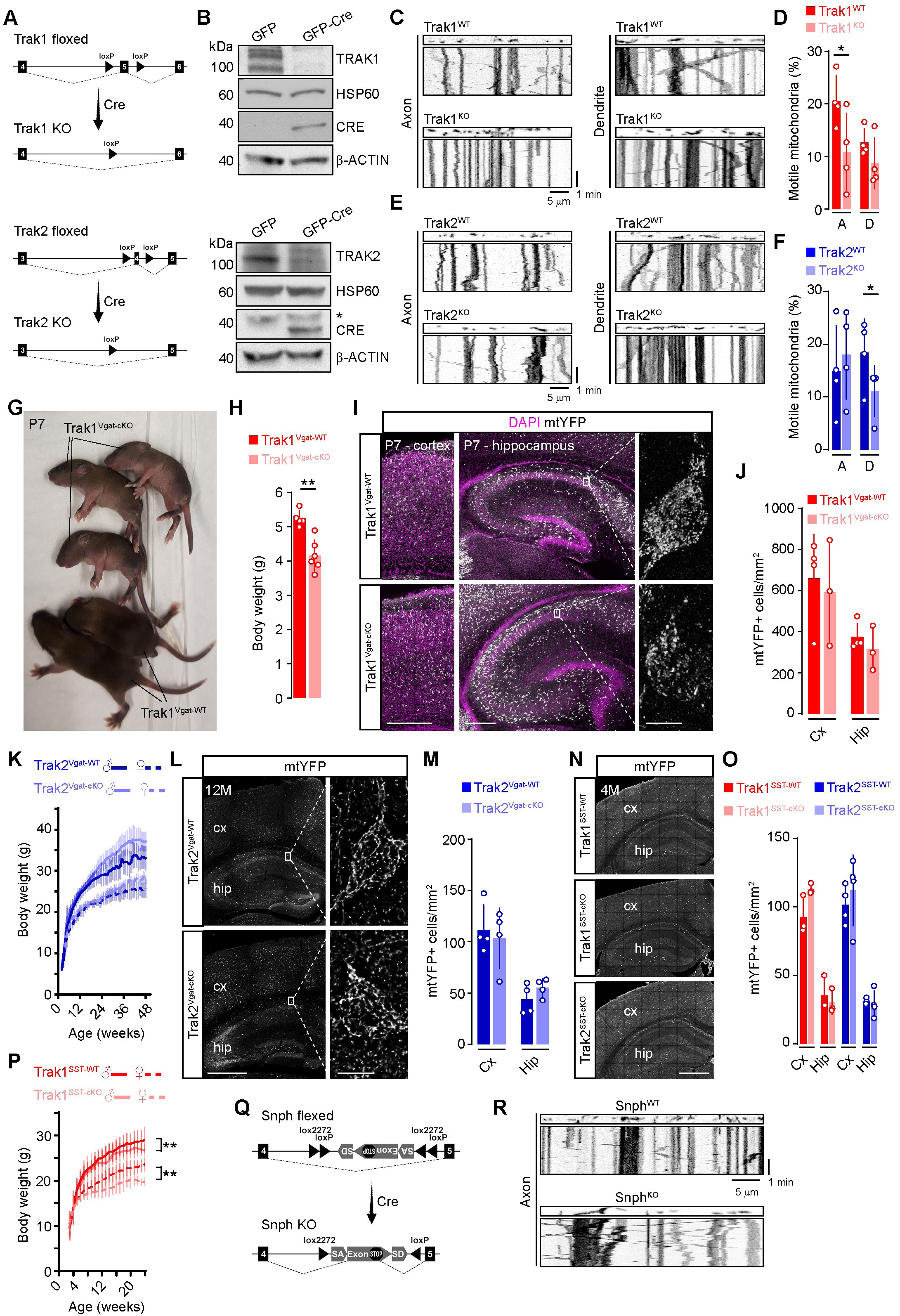
Phenotypic analysis of mice lacking *Trak1* in GABAergic neurons. (**A**) Scheme illustrating the allele targeting strategy flank mouse exon 5 (*Trak1*) and 4 (*Trak2*) with *loxP* sites. **(B)** WB analysis of TRAK1 and TRAK2 expression levels after application of GFP control or GFP-Cre-expressing AAVs in *Trak1* and *Trak2* floxed neurons. *, residual signal of the β-ACTIN antibody. **(C)** Kymograph analysis of mitochondrial motility upon labelling with mitoTracker Red in neuronal axon and dendrites of Trak1^KO^ and control neurons. **(D)** Quantification of mitochondrial motility as shown in C (n= 4 experiments; two-way ANOVA followed by Sidak’s test). A, axon; D, dendrite. **(E)** Kymograph analysis of mitochondrial motility upon labelling with mitoTracker Red in neuronal axon and dendrites of Trak2^KO^ and control neurons. **(F)** Quantification of mitochondrial motility as shown in E (n= 4 experiments; two-way ANOVA followed by Sidak’s test). A, axon; D, dendrite. **(G)** Pictures showing Trak1^Vgat-KO^ and control pups alive at P7. **(H)** Quantification of body weight in Trak1^Vgat-WT^ and Trak1^Vgat-KO^ mice at P6-P7 (n= 5-6 pups per group; unpaired t-test). **(I)** Examples of Trak1^Vgat-WT^ and Trak1^Vgat-KO^ brain sections at P7 expressing mtYFP. Pictures illustrate somatosensory cortex, hippocampus and zooms of neurons in CA1. Bars, 300, 100 and 8 µm. **(J)** Quantification of mtYFP+ cell density in Trak1^Vgat-WT^ and Trak1^Vgat-KO^ mice at P7 (neocortex and hippocampus; n= 3-4 pups per group; two-way ANOVA followed by Sidak’s test). **(K)** Growth curve of Trak2^Vgat-WT^ and Trak2^Vgat-KO^ mice up to 48 weeks of age (n= 2-9 mice per time point and group; mixed effect model followed by Bonferroni’s test). **(L)** Examples of mtYFP+ neurons in hippocampal CA1 of Trak2^Vgat-WT^ and Trak2^Vgat-KO^ mice at 12 months. Bars, 400 and 8 µm. **(M)** Quantification of mtYFP+ cell density Trak2^Vgat-WT^ and Trak2^Vgat-KO^ mice at 12months (neocortex and hippocampus; n= 4 mice per group; two-way ANOVA followed by Sidak’s test). **(N)** Examples of Trak1^SST-WT^, Trak1^SST-cKO^ and Trak2^SST-cKO^ brain sections expressing mtYFP at 4 monts. Bar, 400 µm. **(O)** Quantification of cortical and hippocampal mtYFP+ neurons at 4 months in control, Trak1^SST-cKO^ and Trak2^SST-cKO^ mice (n= 3-4 mice per condition; two-way ANOVA followed by Sidak’s test). **(P)** Growth curve of Trak1^SST-WT^ and Trak1^SST-cKO^ mice up to 24 weeks of age (n= 4-32 mice per time point and group; mixed effect model followed by Bonferroni’s test). **(Q)** Scheme illustrating the allele targeting strategy to generate Snph flexed (double floxed-stop) mice by inserting a double floxed stop exon between endogenous exons 4 and 4 of the *Snph* allele. **(R)** Example of axonal kymograph analysis in primary neurons of Snph flex mice after AAV-based expression of Cre or GFP control. *, p < 0.05. Cx, cortex; hip, hippocampus. See also Figure 3.

**Supplementary Figure 5.**
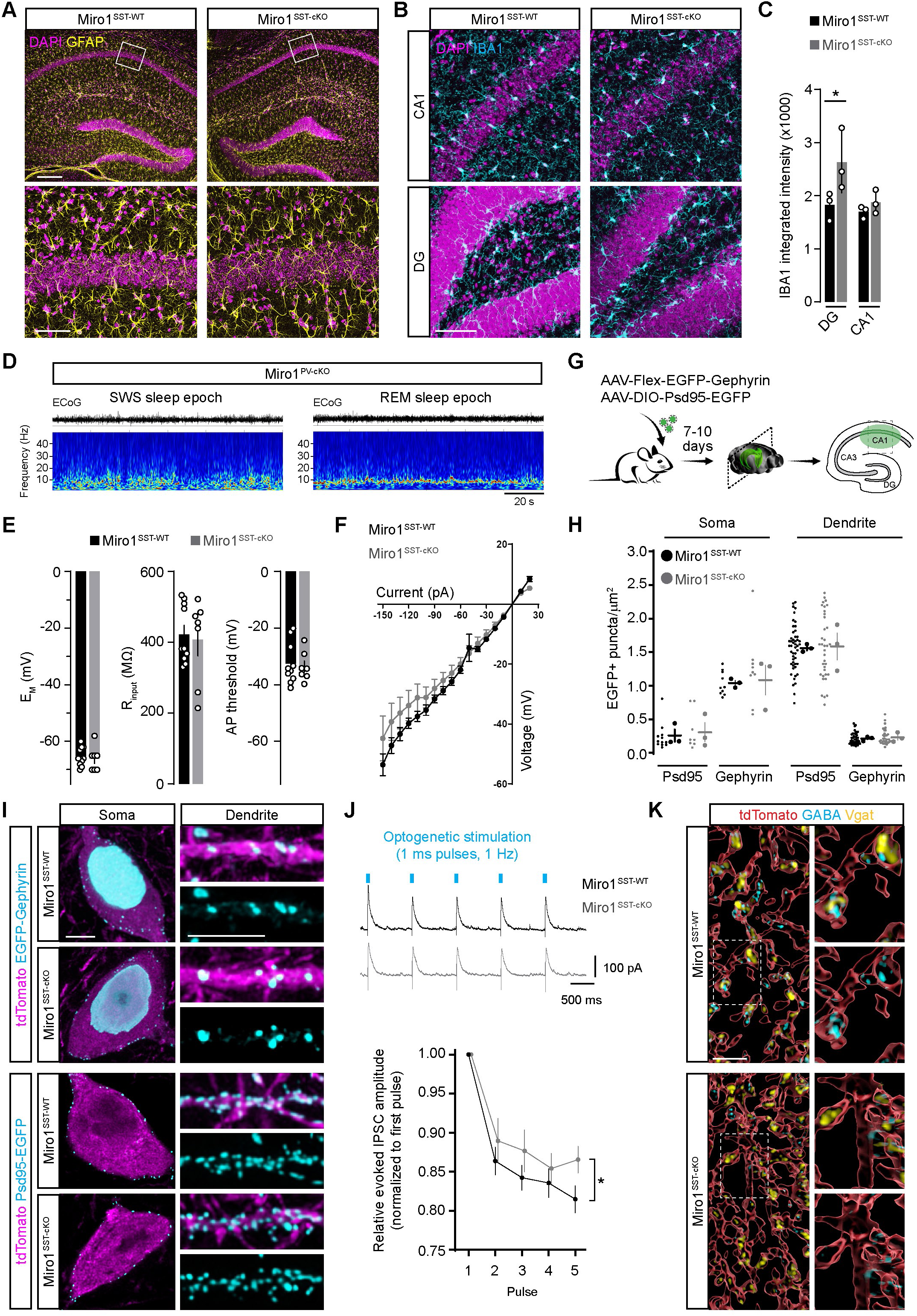
Synaptic changes linked to *Miro1* deletion in SST+ neurons. **(A)** Examples of hippocampal sections immunostained for the astrocytic and microglial markers GFAP and **(B)** IBA1, reporting on their intensity levels in Miro1^SST-WT^ and Miro1^SST-cKO^ mice. Bars, 100 and 60 µm. **(C)** Quantification of IBA1 signal intensity in DG and CA1 as shown in B (n= 3 mice per group; two-way ANOVA followed by Sidak’s test). **(D)** Example of ECoG traces (top) and corresponding wavelet spectrograms (bottom) recorded from Miro1^PV-WT^ and Miro1^PV-cKO^ mice during SWS and REM sleep stages. **(E)** Quantification of resting membrane potential (E_M_), input resistance (R_input_), threshold to action potential and **(F)** current-voltage relationship for CA1 SST tdTomato+ neurons in Miro1^SST-WT^ and Miro1^SST-cKO^ mice (n= 7-11 cells per group obtained from at least 3 mice each; unpaired t-test and Kolmogorov-Smirnov test). **(G)** Scheme showing the used Cre-dependent AAV vectors used to quantify Psd95– and Gephyrin-EGFP. **(H)** Density of EGFP+ puncta in the the cell soma and dendrites of CA1 SST tdTomato+ neurons as shown in I (n= 3 mice per condition, for a total of 8-12 cells and 33-49 dendritic segments; Kruskal-Wallis followed by Dunn’s test). **(I)** Representative examples of cell body and dendrites in AAV-transduced SST+ neurons of Miro1^SST-WT^ and Miro1^SST-cKO^ mice showing the distribution of Psd95– and Gephyrin-EGFP puncta over cytosolic tdTomato. Bars, 5 µm. **(J)** Examples of recordings of pyramidal cells held at 0 mV during 1 Hz optogenetic stimulation of acute slices from Miro1^SST-WT^ and Miro1^SST-cKO^ mice expressing ChR2. Lower graph depicts the relative IPSC amplitude evoked during 5 pulses of stimulation (n= 13-16 cells from at least 3 mice per group; two-way ANOVA followed by Sidak’s test). **(K)** Example of reconstructed portions of axon terminals in the *stratum lacunosum-moleculare* of Miro1^SST-WT^ and Miro1^SST-cKO^ mice, immunostained against Vgat and GABA. Bar, 3 µm. *, p < 0.05. See also Figure 4 and 5.

**Supplementary Figure 6.**
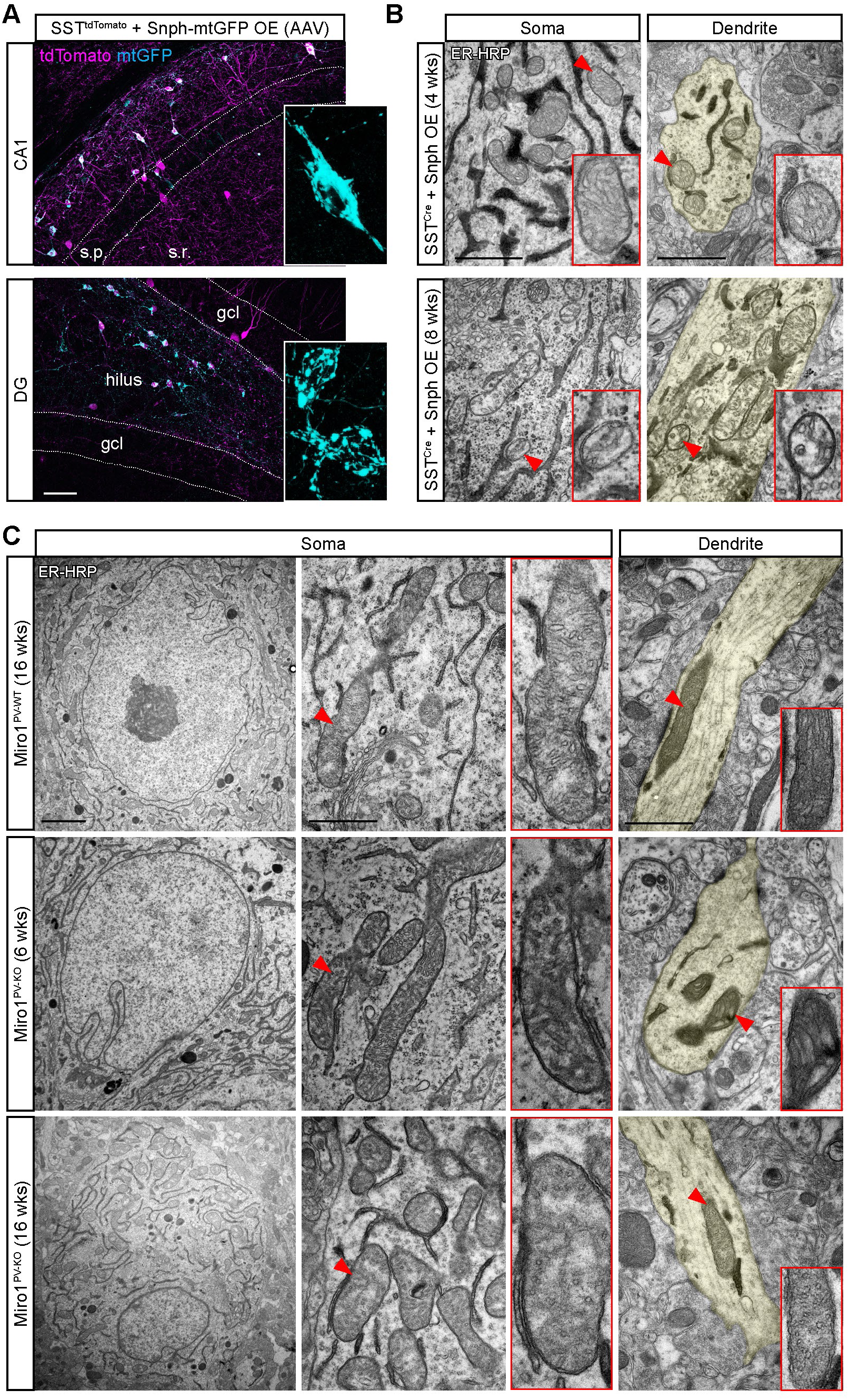
Ultrastructural mitochondrial changes caused by disrupted transport in hippocampal SST+ neurons. **(A)** Examples of tdTomato+ SST neurons in CA1 and DG after ectopic Cre-dependent expression of SNPH and mtGFP. Insets depict zooms of individual cell bodies. Bars, 40 µm. **(B)** Electron micrographs of somatic and dendritic regions of SST neurons labelled with ER-HRP and overexpressing SNPH at 4 and 8 weeks after AAV delivery. Insets show zooms of selected mitochondria (red arrowheads). Bars, 1 µm. **(C)** Electron micrographs of somatic and dendritic regions of PV neurons in Miro1^PV-WT^ and Miro1^PV-cKO^ mice labelled with ER-HRP at 6 and 16 weeks of age. Insets show zooms of selected mitochondria for each region (red arrowheads). Bars, 1 µm. See also Figure 6.

**Supplementary Figure 7.**
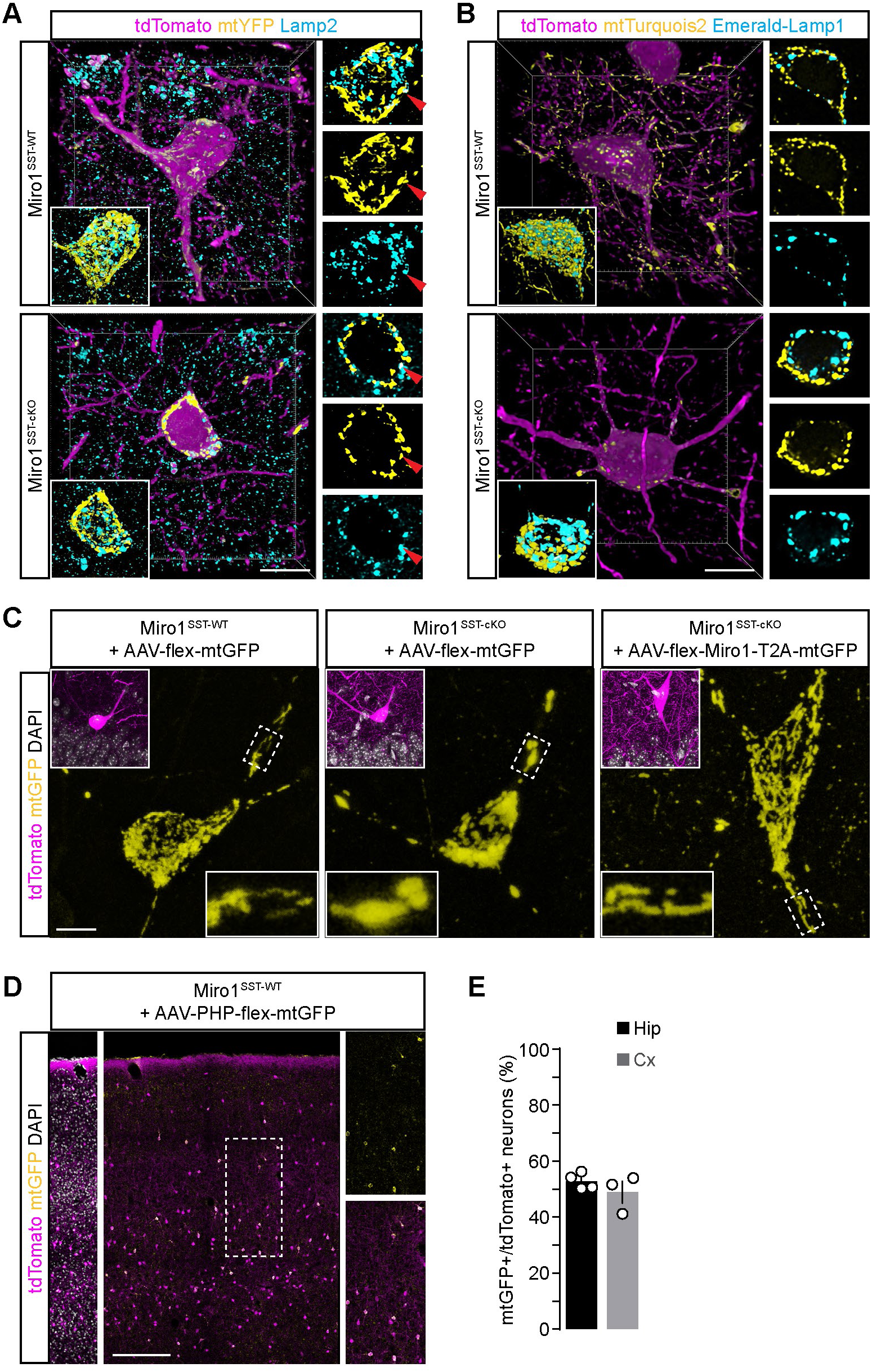
Ultrastructural mitochondrial changes caused by disrupted transport in hippocampal SST+ neurons. **(A)** Representative volumetric reconstructions of mtYFP/tdTomato-expressing SST neurons in the CA1 of Miro1^SST-WT^ and Miro1^SST-cKO^ mice after immunostaining against Lamp2. Lateral panels depict individual and merged channels of the soma within a single optical section, with indicated co-localizing mtYFP/Lamp2 signals (arrowhead). Bar, 5 µm. **(B)** Representative volumetric reconstructions of tdTomato-expressing SST neurons in the CA1 of Miro1^SST-WT^ and Miro1^SST-cKO^ mice after AAV-based co-expression of mtTurquois2 and Emerald-Lamp1. Lateral panels depict individual and merged channels of the soma within a single optical section. Bar, 5 µm. **(C)** Examples of tdTomato-expressing SST neurons in the CA1 of Miro1^SST-WT^ and Miro1^SST-cKO^ mice after AAV-based expression of mtGFP alone or Miro1-T2A-mtGFP. Insets report on zooms of boxed dendritic segments in each image. Bar, 5 µm. **(D)** Overviews and zooms of a portion of the somatosensory cortex in Miro1^SST-WT^ mice after systemic delivery of a Cre-dependent PHP.eB AAV expressing mtGFP, depicting merged and single channels. Bar, 5 µm. **(E)** Quantification of transduction efficiency after systemic delivery of a Cre-dependent PHP.eB AAV expressing mtGFP (n= 3-4 mice; unpaired Mann-Whitney test). See also Figures 6 and 7.

## STAR methods

### Resource availability

#### Lead contact

Further information and requests for resources and reagents should be directed to and will be fulfilled by the Lead Contact, Matteo Bergami.

#### Materials availability

All unique/stable reagents generated in this study are available from the Lead Contact without restrictions. There are restrictions to the availability of mice due to MTA.

#### Data and code availability

All data reported in this paper will be shared by the lead contact upon request. This paper does not report original code. Any additional information required to reanalyze the data reported in this paper is available from the lead contact upon request.

### Experimental model and participant details

Animals were maintained on a C57BL/6N background strain and wild-type as well as transgenic mice of mixed sexes were used in this study. Mice were housed in groups of up to 5 animals per cage supplied with standard pellet food and water *ad libitum* with a 12 h light/dark cycle, while temperature was controlled to 21-22°C. Environmental enrichment (EE) was provided for up 2 days by housing mice in larger “hamster” cages (about 76 × 44 cm) equipped with plastic tunnels and other toys, nesting material and running wheels (Bergami et al., 2015). Thy1^GFP^ mice (line M)(Feng et al., 2000) and mouse lines expressing CRE (SST^CRE^ and PV^CRE^)(Hippenmeyer et al., 2005; Taniguchi et al., 2011) were used as such or crossed with the inducible tdTomato reporter line Ai14 (tdTomato^LSL^)(Madisen et al., 2010) to generate SST^tdTomato^ and PV^tdTomato^ mice. Mice carrying the loxP-flanked genes *Miro1*, *Trak1*, *Trak2* and the double-floxed-stop *Snph* were either used as such or crossed with the Cre-expressing lines SST^CRE^, PV^CRE^, Vgat^CRE^ (Vong et al., 2011) as described in the text. The resulting mice were further crossed with the Ai14 line and/or the mitochondrial-targeted mtYFP^LSL^ inducible reporter line (Sterky et al., 2011), as indicated in the text. All experimental and control animals bearing *Cre* alleles were exclusively maintained in heterozygosity. While growth curves of transgenic lines were examined and reported separately for each sex, no major influence/association of sex on the findings was observed. All experimental procedures were performed in agreement with the European Union and German guidelines and were approved by the State Government of North Rhine Westphalia.

### Method details

#### Mouse genome editing

*Trak1* floxed (C57BL/6N-Trak1^em389Cecad^/Cecad), *Trak2* floxed (C57BL/6N-Trak2^em362Cecad^/Cecad) and Snph flexed (C57BL/6N-Snph^em393Cecad^/Cecad) were generated by CRISPR/Cas9 genome editing. Therefore, a critical exon was identified in *Trak1* (ENSMUSG00000032536) and *Trak2* (ENSMUSG00000026028), which was shared among all annotated full-length transcripts. We classified an exon as critical if deleting it was expected to cause a shift in the reading frame and to create a premature stop codon located at least 55 nucleotides away from the last splice junction. Such changes are likely to trigger nonsense-mediated decay, resulting in the degradation of the mRNA and preventing the production of a functional protein. We identified exon 5 of Ensembl transcript ENSMUST00000045903 for *Trak1* and exon 4 of transcript ENSMUST00000027186 for *Trak2* to be flanked by *loxP* sites for use in Cre-mediated conditional knockout lines. Relevant *Snph* transcripts did not contain any critical exons (ENSMUSG00000027457). We therefore adapted well established FLEx technology (Schnutgen et al., 2003) utilizing a Cre-dependent stable inversion of an artificial stop codon-containing exon into intron 4 of *Snph* transcript ENSMUST00000028951. The artificial exon was composed of an inverted splice-acceptor from the adenovirus major late transcript and *Ifitm2* derived splice-donor flanked by *lox2272* and *loxP* sites. 400 nM of each Alt-R^TM^ guide RNA and 10 ng/µl single-stranded DNA repair template were injected with 200 nM Alt-R^TM^ SpCas9 protein and 30 ng/µl SpCas9 mRNA (TriLink, L-6125-20) in C57BL/6NRj zygotes. All relevant sequences are listed in Supplemental Table 1. CRISPR reagents were purchased from Integrated DNA Technologies (USA) except the Snph FLEx DNA repair template (Genewiz, USA) and animals obtained from Janvier Labs (France). Embryos were cultured *in vitro* and transferred to recipients as described (Wigger et al., 2023). Correct integration of the mutation in the resulting F_0_ mice was validated by Sanger sequencing of a purified PCR amplicon in the founder as well as the F_1_ mice. Genome editing was performed at the *in vivo* Research Facility of the CECAD Research Center, University of Cologne, Germany.

#### Stereotactic procedures and viral injections

Mice were anesthetized by intraperitoneal injection of a ketamine/xylazine mixture (100 mg/kg body weight ketamine, 10 mg/kg body weight xylazine), treated subcutaneously with Carprofen (5 mg/kg) and fixed in a stereotactic frame provided with a heating pad. A portion of the skull covering the somatosensory cortex (from Bregma: caudal: –2.0; lateral: 1.5) was thinned with a dental drill to avoid disturbing the underlying vasculature and a small craniotomy sufficient to allow penetration of a glass capillary was performed. For virus injection a finely pulled glass capillary was then inserted through the dura (from Bregma: –1.3 to –1.2 for CA1; –1.9 to –1.8 for DG) and a total volume of about 300 nl was slowly infused via a manual syringe (Narishige) in two vertical steps spaced by 50 µm each during a time window of 10-20 minutes. After infusion, the capillary was left in place for few additional minutes to allow complete diffusion of the virus. After capillary removal, the scalp was sutured and mice were placed on a warm heating pad until full recovery. The welfare and physical conditions of the animals were monitored daily throughout the duration of the experiment. For cranial window implantation, anesthetized mice received a pre-emptive subcutaneous injection with Carprofen (5 mg/kg) and dexamethasone (0.25 mg/kg). The scalp was removed and the underlying connective tissue was cleared from the skull. A circular craniotomy (3 mm in diameter) was performed over the somatosensory cortex using a dental drill and avoiding to disturb the underlying vasculature. During the whole procedure, a saline solution was flushed onto the area exposed with the craniotomy. A sterile 3 mm circular glass coverslip (#1 thickness, Warner Instruments) was gently implanted into the craniotomy site and sealed in place with a thin layer of Sylgard (Sigma) before applying dental cement (Dentalon plus, Heraeus Kulzer GmbH) to fix the coverslip and cover the surrounding exposed skull. An aluminium chamber plate (CP-1, Narishige) was fixed with cement on top of the cover glass to facilitate mouse head immobilization at the 2-photon microscope via a head holder (MAG-2, Narishige). The depth of anaesthesia was assessed throughout the surgery and recording time (usually 1-2 hours) and eventually mice received one or more additional boluses of anaesthetic each corresponding to one third of the initial dose.

#### Viral production

Helper-free AAV vectors were either obtained from Addgene or produced according to adapted established methods. Both variants used, AAV1 (for intracranial delivery) and AAV-PHP.eB (for systemic delivery), were produced in 293AAV cells (Cell Biolabs; AAV-100) by transient transfection with plasmid containing desired transgene, helper plasmid and variant-specific packaging Rep/Cap plasmid. Viral particles were harvested, purified by discontinuous iodixanol gradient ultracentrifugation (24h at 32,000 rpm and 4°C) and finally concentrated using Amicon® Ultra Centrifugal Filter unit with a 100 kDa molecular weight cut-off. Virus titers were measured by determining the number of DNAse I-resistant viral genome (vg) using quantitative PCR (qPCR), and viral preparations stored at –80°C until use.

#### In vivo 2-photon imaging

In vivo imaging was conducted using a multiphoton microscope (TCS SP8 MP-OPO, Leica Microsystems). The setup included a Chameleon Vision II laser (wavelength range: 689-1040 nm), a water-immersion IR Apo L25x/0.95 W objective, and two internal HyD detectors with specific filter settings (FITC: 500-550 nm, TRITC: 565-605 nm). An adjustable Scientifica stage was used, and the anesthetized mouse laying on a warm pad heated at 37 degrees was secured to a head device (MAG-2, Narishige), which was connected to the imaging stage. During the imaging sessions, the mouse was monitored using an infrared camera. The laser was tuned to 900 nm to simultaneously visualize tdTomato and mtYFP signals. The imaging acquisition rate was set at one frame every 2 seconds for a duration of 3 minutes.

#### Acute brain slice preparation

Freshly removed mouse brains were placed in ice-cold, carbogen-saturated (5% CO2, 95% O2) artificial cerebrospinal fluid (aCSF) containing (in mM): 125 NaCl, 2.5 KCl, 1.25 NaH2PO4, 25 NaHCO3, 25 D-Glucose, 0.5 CaCl2 and 3.5 MgCl2, adjusted to pH 7.4 and 310-320 mOsm. Coronal hippocampal slices (300 μm thick) were prepared with a vibratome (Micron HM650V ThermoScientific, Walldorf, Germany; or VT1200 S, Leica Biosystems GmbH, Nussloch, Germany) in ice-cold aCSF. The obtained slices were transferred into a chamber containing a continuously carbogen-saturated aCSF at room temperature with the following composition (in mM): 125.0 NaCl, 2.5 KCl, 1.25 NaH2PO4, 25.0 NaHCO3, 25.0 D-Glucose, 2.5 CaCl2, 1.3 MgCl2, adjusted to pH 7.4 and 310-320 mOsm. For experiments involving recordings of pyramidal neurons in CA1 (including optogenetic experiments), the used carbogen-saturated (5% CO2, 95% O2) aCSF contained (in mM): 130 NaCl, 24 NaHCO_3_, 3.5 KCl, 1.25 NaH_2_PO_4_, 1 CaCl_2_, 2 MgCl_2_, 10 glucose (pH 7.4). Slices were stored for at least 30 min to allow tissue recovery before starting recordings or imaging sessions.

#### Live imaging and analysis of mitochondrial movement ex vivo and in culture

For *ex vivo* experiments, acute slices were moved into a dedicated imaging chamber and experiments were conducted under continuous aCSF perfusion (2 ml/min with recording solution) at a constant temperature of 32-33°. Imaging was performed using a multiphoton laser-scanning microscope (TCS SP8 MP-OPO, Leica Microsystems) equipped with a Leica 25x objective (NA 0.95, water) and a Ti:Sapphire laser (Chameleon Vision II, Coherent). For mitochondrial transport analysis, the excitation wavelength was set at 900 nm, and the rate of image acquisition was one frame every 10 s for up to 5 min. Imaging was conducted at least 20-30 μm below the slice surface, focusing on mtYFP+ cell bodies or axonal segments in the *stratum radiatum* or *oriens* of the CA1 region. Only slices with a general healthy appearance throughout the recording time and whose acquisitions displayed only no or only minor spatial drift in z during the whole imaging session were included in subsequent analysis. Time-lapse image sequences were drift-corrected by utilizing the “fast & rigid body” options of the TurboReg plugin (http://bigwww.epfl.ch/thevenaz/turboreg/) in ImageJ, aligning each frame to a median projection of eleven frames centered on the middle of the time series. In case of non-satisfactory results, the moco plugin (https://github.com/NTCColumbia/moco) was used alternatively. For analysis of mitochondrial trafficking in neuronal cultures, cells were first incubated for 10 min in 10nM MitoTracker™ Red CMXRos (M7512, Invitrogen) in Neurobasal and then washed three times with HEPES-buffered Tyrode’s solution. Imaging was performed in HEPES-buffered Tyrode’s solution (119 mM NaCl, 5 mM KCl, 25mM HEPES buffer, 2mM CaCl2, 2 mM MgCl2, 33mM glucose) maintained at 37°C using a 40x (1.3NA) oil objective on a TCS SP8 gSTED. Videos were acquired at 0.5 frames per second for 5 min. For quantification of mitochondrial motility in culture, *ex vivo* and *in vivo*, kymographs were generated using neuronal processes linearization via ImageJ (Straighten function) followed by Multiple Kymograph plugin. Mitochondrial movement was assessed by quantifying the percentage of motile mitochondria (at least 2 µm displacement) during the 3-5 min recording time, normalized by the total number of visible mitochondria at the first frame of the movie.

For analysis of mitochondrial membrane potential *ex vivo*, slices were first incubated for 30 min in 100nM tetramethylrhodamine methyl ester (TMRM, Thermo Fisher Scientific) added to aCSF at RT. TMRM and mtYFP signals were imaged by using an excitation wavelength of 920 nm and TRITC (585/40)/FITC (525/50) filters, performing appropriate controls to rule out signal bleed-through. Acquisition was performed through z-stacks spanning first 50 μm of the slice thickness, followed by removal from analysis of the first 20 μm to rule out potential damage of the slice surface. Quantification was performed by masking the area of individual mtYFP+ neurons in single stacks and normalizing the signal of TMRM onto mtYFP in the same cells.

#### Electrophysiology

Whole-cell patch-clamp recordings of SST+ neurons were performed with an electrophysiology rig (Scientifica SliceScope Pro, U.K.) under visual control with a fixed stage upright microscope equipped with 40x water-immersion objective. Neurons were visualized and identified for their expression of tdTomato and their anatomical positioning in the CA1 *stratum oriens* by means of differential-interference-contrast (DIC)-infrared optics and epifluorescence along with a CCD camera (Orca-ER, Hamamatsu, Shizouka, Japan). Patch pipettes with a tip resistance of 5-10 MΩ were made from borosilicate glass capillaries (0.86 mm ID, 1.5 mm OD, GB150-8P, Science Products, Hofheim, Germany) with a horizontal pipette puller (Model P-1000, Sutter Instruments, Novato, CA, USA). Immediately before recordings, the patch pipette was filled with internal solution containing, in mM: 4.0 KCl, 2.0 NaCl, 0.2 EGTA, 125.0 K-Gluconate, 10.0 HEPES, 4.0 ATP(Mg), 0.5 GTP(Na), 10.0 phosphocreatin, adjusted pH 7.25 and 290 mOsm. Whole-cell patch clamp recordings were performed with an ELC-03XS patch clamp amplifier (npi electronic GmbH, Tamm, Germany) and a data acquisition unit (Micro1401-4, Science products Gmbh, Germany) controlled by the software Signal (version 6.0, Cambridge Electronic, Cambridge, UK). The data were sampled with a frequency of 10 kHz and filtered by two short-pass Bessel filters set at 3 kHz for voltage clamp (VC) and 12 kHz for current clamp (CC). Offset potentials and capacitive currents were compensated manually. Whole-cell capacitance was determined by using the capacitance compensation (Cslow) of the amplifier. Shortly after establishing whole-cell configuration in voltage-clamp mode, the resting membrane potential (E_M_) was measured by applying 0 pA. To determine basic electrophysiological properties, different current injection protocols were executed during the recordings. First, spontaneous postsynaptic currents (sPSCs) were recorded for 5 min in VC mode with holding potential set to –70 mV in in absence of external stimulation. Frequency and amplitude of single current events were quantified by using a customized script. To analyze passive membrane properties (input resistance and membrane time constant), three hyperpolarizing current pulses of –5 pA increments with a duration of 1 s were applied. To analyze action potential properties, depolarizing current pulses starting from +20 with a duration of 2 s were injected. The number of sweeps was individually adjusted for each cell until maximum spike frequency was reached. Calculation of action potential threshold was obtained from the second derivative of the first action potential triggered by our protocol. Finally, an I/V curve was performed by applying 2 s long current injections from –150 pA to +40 pA with +10 pA increments. Data of electrophysiological recordings were analyzed with Igor Pro (version 7.05.2, WaveMetrics, USA).

Whole-cell patch-clamp recordings of CA1 pyramidal neurons was performed using a LN Scope (Luigs&Neumann GmbH, Ratingen Germany) with infrared Dodt gradient contrast illumination and a 40x water-immersion objective (Olympus Deutschland GmbH, Hamburg, Germany) connected to Photometrics Moment sCMOS camera (Photometrics, Huntington Beach, CA, USA). Hippocampal CA1 pyramidal neurons were recognized by their typical location in the stratum pyramidale and absence of tdTomato fluorescence. Patch-clamp pipettes were pulled from 3.3 borosilicate glass capillaries (Hilgenberg GmbH, Malsfeld, Germany) and had a resistance of 3-5MΩ when filled with the following intracellular solution (in mM): 125 Cs-methanesulfonate, 0.3 GTP-Na, 4 ATP-Mg_2_, 16 KHCO_3_, 10 QX-314-Cl (pH7.3). In some recordings, biocytin (0.3 – 0.5%) was added to the solution to characterize the morphology of the recorded cell after fixation. Recordings were performed using a Multiclamp 700B amplifier (Molecular Devices, San Jose, CA). Signals were filtered at 3 kHz and digitized at 20 or 50 kHz using a Digidata 1550B equipped with a Hum Silencer and the Clampex 11 program suite (Molecular Devices). Gap free voltage clamp recordings of sEPSCs and sIPSCs were performed at the holding potential indicated in the text and/or legends (–60 mV and 0 mV, respectively) for 3 minutes. For optogenetic experiments, light stimulation was achieved by a Solis 470 LED (Thorlabs GmbH, Mölndal, Germany) attached to the epifluorescence port of the microscope. Brief light flashes (1 ms, 50% power) were directed to the slice in proximity of the recorded neuron. Stimulation was repeated for at least three times with 60s between each repetition. Photoevoked IPSCs were recorded at a holding potential of 0mV. The events were analysed using the Clampfit 11 program suite (Molecular Devices) utilizing template search for sPSC frequency and peak amplitude measurements after light stimulation for photoevoked IPSCs.

#### Electrocorticogram recordings

Telemetric electrocorticogram analyses were performed by using implantable radio transmitters (models TA11EA-F10 or TA11ETA-F10, DataSciences International). Mice received 0.05=mg/kg buprenorphine (subcutaneously) for analgesia and were anesthetized with 1.5–2% isoflurane in 100% oxygen. Midline skin incisions were made on the top of the skull and in the neck. The transmitter body was implanted subcutaneously. The tips of the leads were placed 2=mm posterior to bregma and 1.8=mm right to the midline for the recording electrode, and 1=mm posterior to lambda and 1–2=mm left to the midline for the reference electrode. The electrodes were fixed with dental cement. Radio transmitters allowed simultaneous monitoring of electrocorticogram and motor activity in undisturbed, freely moving mice housed in their home cages. Telemetry data and corresponding video data were recorded 48–72=h after surgery and were stored digitally using Ponemah software (Data Sciences International). Electrocorticogram recordings lasted over 50=h, sampled at 500=Hz, with continuous video recording.

#### Doxycycline treatments

For treatments in culture, single or multiple pulses of Dox (Sigma, 100 ng/mL) in Neurobasal were applied for 30 min each, followed by a washing step and supplementation of Dox-free complete Neurobasal until the end of the experiment. For experiments in vivo, about 10 days after AAV delivery mice were injected intraperitoneally with a single or multiple Dox pulses (dissolved in sterile saline) as indicated in the text, each corresponding to 50 µg/g of body weight. For quantification of relative mitochondrial turnover rates in EE-exposed and Miro1^SST-cKO^ mice, the paradigm involved a first pulse 4 days and a second one a day before sacrifice. Both in culture and *in vivo*, validation of lack of leakiness of the system was performed following no Dox treatment or application of vehicle only.

#### Systemic AAV delivery

Mice during their 4th week of life were subjected to standard tail vein injection procedure of a 100μL volume using a 30G needle insulin syringe containing 20μL of virus + 80μL of sterile saline. Used viruses included AAV-PHP.eB-FLEX-mtGFP, AAV-PHP.eB-FLEX-Miro1-T2A-mtGFP, or AAV-PHP.eB-FLEX-Trak1.

#### Transmission electron microscopy

Anesthetized mice were transcardially perfused with a fixative solution containing 2% formaldehyde and 2.5% glutaraldehyde in 0.1 M cacodylate buffer. The brain was isolated, cut in 50-100 μm thick coronal sections using a VT1000 S microtome (Leica). Samples were washed two times with 0.1 M sodium cacodylate buffer, and free aldehyde groups were quenched using 0.1M Glycin in 0.1M Cacodylate buffer for two times 20 min. After a short wash with 0.1M Cacodylate buffer, slices were incubated for 30min in 0.5mg/ml Diaminobenzidine in 0.1M Cacodylate buffer with 0.03% H2O2. Brain slices were washed three times with 0.1M Cacodylate buffer and trimmed to the region of interest using razor blades. Post-fixation was applied using 1% Osmiumtetroxid and 1.5% Potassium hexacyanoferrate for 30 min at 4°C. After 3×5min wash with ddH2O, samples were dehydrated using ascending ethanol series (50%, 70%, 90%, 100%) for 10 min each. Infiltration with a mixture of 50% Epon/ethanol overnight at 4°C and with pure Epon for two times 2 h. Samples were embedded into TAAB capsules and cured for 48 h at 60°C. Ultrathin sections of 70 nm were cut using an ultramicrotome (Leica Microsystems, UC6) and a diamond knife (Diatome, Biel, Switzerland) and stained with 1.5 % uranyl acetate for 15 min at 37°C and Reynolds lead citrate solution for 4 min. Images were acquired using a JEM-2100 Plus Transmission Electron Microscope (JEOL) operating at 80kV equipped with a OneView 4K camera (Gatan). For analysis, electron micrographs of cells, dendrites and axon terminals labelled with HRP or APEX2 were acquired with a digital zoom of 5000x or 10000x.

#### Cell sorting by flow cytometry

Mouse hippocampi were microdissected as previously reported (Motori et al., 2020) and chopped in ice-cold dissociation medium (HBSS Ca^2+^– and Mg^2+^-free, supplemented with 20 mM glucose, penicillin 50 U/ml, and streptomycin 0.05 mg/ml) and then digested in papain medium (HBSS, supplemented with l-cysteine·HCl 1 mg/ml, papain 16 U/ml, and DNase I 0.1 mg/ml) at 37°C for 30 min. Tissues were first washed in HBSS medium containing ovomucoid (10 mg/ml), BSA (10 mg/ml), and DNase (0.1 mg/ml) at room temperature to block the enzymatic digestion and then gently triturated in HBSS containing 20 mM glucose, penicillin (50 U/ml), streptomycin (0.05 mg/ml), and DNase (0.1 mg/ml) to release single cells. The resulting cell suspension was filtered through a 70-μm cell strainer, and then cells were pelleted by centrifugation (1110 rpm, 5 min, 4°C) and resuspended in sorting medium (HBSS, supplemented with 20 mM glucose, 20% fetal bovine serum, penicillin 50 U/ml, and streptomycin 0.05 mg/ml); cell viability was assessed by using propidium iodide, and the cell suspension was filtered through a 50-μm cell strainer before flow cytometry. TdTomato– and GFP-expressing cells in suspension were stained with DAPI and DRAQ5 to identify intact and viable neurons. The DAPI negative/ DRAQ5 positive/ tdTomato positive and the DAPI negative/ DRAQ5 positive/ GFP positive cell populations were selected. Cell sorting was performed using BD FACSAria IIIu and BD FACSAria Fusion with FACSDiva 8.0.1 software. Neurons were sorted at 4°C using a 100 µm nozzle and sheath pressure was set at 20 psi. 0.9% NaCl was used as sheath fluid. Collected cells were then pelleted by centrifugation (1110 rpm, 5 min, 4°C) and stored at −80°C until further use.

#### Mass spectrometry (MS)

LCMS Data Independent Acquisition: samples were analyzed by the CECAD Proteomics Facility on a Q Exactive HFx mass spectrometer that was coupled to an EASY nLC 1200 (Both Thermo Scientific). Peptides were loaded with solvent A (0.1% formic acid in water) onto an in-house packed analytical column (50 cm — 75 µm I.D., filled with 2.7 µm Poroshell EC120 C18, Agilent). Peptides were chromatographically separated at a constant flow rate of 250 nL/min and the following gradient: 4-23% solvent B (0.1% formic acid in 80 % acetonitrile) within 65.0 min, 23-55% solvent B within 13.0 min and 55-95% solvent B within 3.0 min, followed by a 9 min column wash with 95% solvent B. The mass spectrometer was operated in data independent acquisition mode (DIA). MS1 scans were acquired from 350 m/z to 1650 m/z at 120k. Maximum injection time was set to 50 msec and the AGC target to 3e6. 24 variable DIA windows covering the precursor range from 350 – 1650 m/z were acquired at 30k resolution with a maximum injection time of 60 msec and an AGC target of 3e6. All scans were stored as centroid.

Data processing and analysis: the raw data files were processed using Spectronaut 19 (Biognosys) in direct DIA mode with a Swissprot mouse canonical fasta file (UP000000589, downloaded 15/01/2024). Cross run normalization was enabled and differing from the default settings stripped sequence quantities were calculated as summed intensities of precursors ions. MaxLFQ was used for protein quantification using proteotypic stripped sequences only. The different mouse lines used in this study (THY1-GFP, PAR^tdTomato^, SOM^tdTomato^) were processed separately. Normalized protein group quantities together with iBAQ values, number of stripped sequences and the corresponding protein and gene identifiers from Spectronaut pivot reports were imported into Perseus 1.6.15 (Tyanova, 2016) for further statistical analysis. The dataset was filtered for at least 100 % data completeness in at least one biological condition and at least 3 stripped sequences per protein experiment wide. Missing values were imputed from the left end of the normal distribution using Perseus default settings. Student’s T-test was used to test for significant (FDR<0.05, S0=0.2) changes in protein abundance across the biological conditions. The data set was annotated with MitoCarta 3.0, GO, KEGG as well as Reactome terms. Relative enrichment of ontology terms in the group of significantly changed proteins was determined using the Fisher exact test. In addition, 1D enrichment analysis were calculated on the Log2 T-test differences.

#### Metabolomics

Dissected brains were maintained in cold PBS 1x and hippocampus was taken out. The tissue was then rinsed three times with 75 mM Ammonium carbonate (pH= 7.4) and then placed on aluminium foil on dry ice. Frozen samples were processed by the metabolomics facility of Max Planck Institute for Biology of Ageing (Cologne, Germany).

Anion-Exchange Chromatography Mass Spectrometry (AEX-MS) for the analysis of anionic metabolites: the polar phase of the extracted metabolites was re-suspended in 400 µl of ULC/MS grade water (Biosove, Valkenswaard, Netherlands). After 15 min incubation on a thermomixer at 4°C and a 5 min centrifugation at 16.000 x g at 4°C, 100 µl of the cleared supernatant were transferred to polypropylene autosampler vials (Chromatography Accessories Trott, Germany). The samples were analysed using a Dionex ionchromatography system (Integrion, Thermo Fisher Scientific) as described previously 1. In brief, 5 µL of polar metabolite extract were injected in full loop mode using an overfill factor of 1, onto a Dionex IonPac AS11-HC column (2 mm × 250 mm, 4 μm particle size, Thermo Fisher Scientific) equipped with a Dionex IonPac AG11-HC guard column (2 mm × 50 mm, 4 μm, Thermo Fisher Scientific). The column temperature was held at 30°C, while the auto sampler was set to 6°C. A potassium hydroxide gradient was generated using a potassium hydroxide cartridge (Eluent Generator, Thermo Scientific), which was supplied with deionized water. The metabolite separation was carried at a flow rate of 380 µL/min, applying the following gradient conditions: 0-3 min, 10 mM KOH; 3-12 min, 10−50 mM KOH; 12-19 min, 50-100 mM KOH, 19-21 min, 100 mM KOH, 21-22 min, 100-10 mM KOH. The column was re-equilibrated at 10 mM for 4 min. For the analysis of metabolic pool sizes the eluting compounds were detected in negative ion mode using full scan measurements in the mass range m/z 50 – 750 on a Q-Exactive HF high resolution MS (Thermo Fisher Scientific). The heated electrospray ionization (ESI) source settings of the mass spectrometer were: Spray voltage 3.2 kV, capillary temperature was set to 300°C, sheath gas flow 50 AU, aux gas flow 20 AU at a temperature of 330°C and a sweep gas glow of 2 AU. The S-lens was set to a value of 60. The semi-targeted LC-MS data analysis was performed using the TraceFinder software (Version 4.1, Thermo Fisher Scientific). The identity of each compound was validated by authentic reference compounds, which were measured at the beginning and the end of the sequence. For data analysis the area of the deprotonated [M-H+]-1 or [M-2H]-2 monoisotopic mass peak of each compound was extracted and integrated using a mass accuracy <5 ppm and a retention time (RT) tolerance of <0.05 min as compared to the independently measured reference compounds. Areas of the cellular pool sizes were normalized to the internal standards, which were added to the extraction buffer, followed by a normalization to the fresh weight of the analyzed sample.

Semi-targeted liquid chromatography-high-resolution mass spectrometry-based (LC-HRS-MS) analysis of amine-containing metabolites: 50 µL of the available 400 µL of the above mentioned (AEX-MS) polar phase were mixed with 25 µl of 100 mM sodium carbonate (Sigma), followed by the addition of 25 µl 2% [v/v] benzoylchloride (Sigma) in acetonitrile (UPC/MS-grade, Biosove, Valkenswaard, Netherlands). Derivatized samples were thoroughly mixed and kept at 20°C until analysis. For the LC-HRMS analysis, 1 µl of the derivatized sample was injected onto a 100 x 2.1 mm HSS T3 UPLC column (Waters). The flow rate was set to 400 µl/min using a binary buffer system consisting of buffer A (10 mM ammonium formate (Sigma), 0.15% [v/v] formic acid (Sigma) in UPC-MS-grade water (Biosove, Valkenswaard, Netherlands). Buffer B consisted of acetonitrile (IPC-MS grade, Biosove, Valkenswaard, Netherlands). The column temperature was set to 40°C, while the LC gradient was: 0% B at 0 min, 0-15% B 0– 4.1min; 15-17% B 4.1 – 4.5 min; 17-55% B 4.5-11 min; 55-70% B 11 – 11.5 min, 70-100% B 11.5 – 13 min; B 100% 13 – 14 min; 100-0% B 14 –14.1 min; 0% B 14.1-19 min; 0% B. The mass spectrometer (Q-Exactive Plus, Thermo Fisher Scientific) was operating in positive ionization mode recording the mass range m/z 100-1000. The heated ESI source settings of the mass spectrometer were: Spray voltage 3.5 kV, capillary temperature 300°C, sheath gas flow 60 AU, aux gas flow 20 AU at 330°C and the sweep gas was set to 2 AU. The RF-lens was set to a value of 60. Semi-targeted data analysis for the samples was performed using the TraceFinder software (Version 4.1, Thermo Fisher Scientific). The identity of each compound was validated by authentic reference compounds, which were run before and after every sequence. Peak areas of [M + nBz + H]+ ions were extracted using a mass accuracy (<5 ppm) and a retention time tolerance of <0.05 min. Areas of the cellular pool sizes were normalized to the internal standards (U-15N;U-13C amino acid mix (MSK-A2-1.2), Cambridge Isotope Laboratories), which were added to the extraction buffer, followed by a normalization to the fresh weight of the analyzed sample.

#### Immunohistochemistry

Fixed brains (4% paraformaldehyde in PBS) were washed three times with 1x PBS for 10 minutes each at RT and 50 to 70 µm thick coronal sections (depending on the analysis) were serially cut using a VT1000 S microtome (Leica). Free-floating brain slices were incubated in blocking buffer (0.1 M Phosphate buffer, 0.3% Triton X-100, 5% bovine albumin serum) for 1 h with gentle agitation. Sections were then incubated with primary antibodies diluted in blocking buffer overnight at 4°C. Slices were washed three times in 1x PBS for 10 minutes each at RT and then incubated with secondary antibodies diluted in 3% BSA for 2 hours at RT with gentle agitation. Sections were then rinsed three times in 1x PBS for 10 min each at RT, counterstained with 4’,6-diamidino-2-phenylindole (DAPI, ThermoFisher, 3 μM) in the last washing step and mounted on microscopic slides with Aqua Poly/Mount (Polysciences). The following primary antibodies were used for immunohistochemistry: chicken anti-GFP (1:500, Aves Labs, GFP-1020), rabbit anti-RFP (1:500, Rockland, 600-401-379), mouse anti-GFAP (1:1000, Millipore, MAB360), rabbit anti-Iba1 (1:1000, Wako, 019-19741), guinea pig anti-NeuN (1:1000, Millipore, ABN90P), rat anti-Somatostatin (1:500, Millipore, MAB354), guinea pig anti-Parvalbumin (1:800, Synaptic Systems, 195004), mouse anti-ATP5B (1:500, Abcam, AB5432), mouse anti-MTCO1 (1:500, Abcam, AB14705), rabbit anti-TOM20 (1:500, Santa Cruz, SC-11415), rabbit anti-RHOT1 (1:50, Atlas antibodies, HPA010687), mouse anti-c-fos (1:500, Abcam, ab7963), guinea pig anti-Vgat (1:500, Synaptic Systems, 131004), rat anti-Lamp2 (1:500, Abcam, ab13524). The following secondary antibodies (donkey) were used: Alexa Fluor 488-, Alexa Fluor 546-, Alexa Fluor 647– conjugated secondary antibodies anti rabbit, mouse, chicken, rat and guinea pig (1:1000, Jackson ImmunoResearch, 703545155/ 703605155/ 711605152/ 712605150/ 705606147/ 715605150/ 706605148). Images (1024 x 1024 or 512×512) were acquired on a confocal laser scanning microscope (TCS SP8 gSTED 3x, Leica Microsystems) equipped with following objectives: Apo 20x/0,75 Multi-Immersion, PL Apo 40x/1,3 Oil DIC, PL Apo 63x/1,2 water DIC, PL Apo 100x/1.40 Oil STED Orange objectives. Following lasers were used: 405 nm, 458 nm, 477 nm, 488 nm, 514 nm, white light laser (470-670 nm tunable).

#### Primary neuronal cultures

Brain embryos (IZlE13.5) were isolated following cervical dislocation of the anesthetized pregnant mother. Dissected cortices were collected in cold Hanks’ balanced salt dissociation solution (HBSS) and dissociated in pre-warmed Dulbecco’s modified Eagle’s medium (DMEM) containing papain (2.5 U/ml, Sigma) at 37°C for 15 min. Medium was then replaced by DMEM with 10% fetal bovine serum (FBS) and mechanically triturated. Cells were pelleted by centrifugation for 5 min at 1100 rpm at 4°C and then pre-plated in 60 mm culture dish at 37°C for 20 min. Supernatant was collected and cell count was determined with an automated cell counter (Invitrogen, #A50298). Neurons were then plated at the desired density in Neurobasal serum-free medium containing 1% B27 supplement and 0.5 mM GlutaMax and maintained at 37°C and 5% CO_2_ throughout the experiment. For viral infection, 30% of medium was first removed from wells, the virus added to the remaining volume, and after 4 h complete neurobasal medium was added to reach initial volume.

#### Western Blotting

Neurons were placed on ice and washed once with 1x PBS, then scraped off in 2 ml ice-cold 1x PBS and centrifuged for 5 min at 2000 rpm. Pellet was resuspended in cold lysis buffer (0.5% Triton x-100, 0.5% Deoxygenate and 1/4 tablet Protease Inhibitor, Roche Diagnostics), followed by 1 h incubation on ice. Protein concentration was determined by Bradford assay. Desired amount of protein lysate was suspended in Laemmli sample buffer and incubated for 5 min at 95 °C. Proteins were then separated by SDS–polyacrylamide gel electrophoresis and blotted onto methanol activated polyvinylidene fluoride membranes (Amersham, GE Healthcare). Following blocking, membranes were incubated with primary antibodies diluted in 3% milk powder overnight at 4 °C. The next day, the membranes were washed 2 times with TBST (Tris Buffered Saline with Tween) for 5 min and incubated for 2h at RT in the appropriated horse radish peroxidase (HRP) conjugated secondary antibody in TBST, followed by further washes with TBST. Blots were developed with ECL solution (Amersham, GE Healthcare) according to the manufacturer’s protocol, followed by exposure to x-ray film (Fujifilm Super RX). The following primary antibodies were used: anti-RHOT1 (Sigma, HPA010687, rabbit, 1:500), Anti-HSP60 (Santa Cruz, sc-1052, goat, 1:500), anti-CRE (BioLegend, 908001, rabbit, 1:500), anti-β-3-TUBULIN (Synaptic Systems, 302304, guineapig, 1:1000), anti-TRAK1 (Sigma, HPA005853, rabbit, 1:500), anti-TRAK2 (sigma, HPA015827, rabbit 1:500), anti-β-ACTIN (Sigma, A2228, mouse, 1:1000) and anti-SNPH (Abcam, ab192605, rabbit, 1:1000). The following HRP conjugated secondary antibodies against mouse (Sigma, A9044, 1:5000), rabbit (Sigma, A0545, 1:5000) and goat (Santa Cruz, sc-2384, 1:5000) were used.

#### Image analysis and quantification

For quantification of cell densities, 3-5 sections per mouse brain corresponding to similar anatomical locations across mice were used. All quantifications were done on z-stack confocal images after their conversion into TIFF files, and analysis was performed off-line using the ImageJ software (National Institute of Health, Bethesda, United States). Quantification of cell densities across hippocampus and cortex was performed manually on acquisitions taken with a 20x objective and by using the Cell Counter plugin, by normalizing the number of marker-positive cells over the area or volume of the corresponding region of interest. For high-resolution imaging of mitochondria within single cells and compartments in fixed tissue, mice bearing a mitochondrial reporter, or immunostained for mitochondrial markers, were used. Z-stack acquisitions of individual cell regions of interest were taken with a 63x or 100x oil objectives (and additional digital zooms when required) utilizing an inter-stack interval of 0.3 μm or less and a resolution of 1024×1024 pixels. Single-cell quantification of ATP5B and MT-CO1 expression levels was performed in presence or absence of normalization over TOMM20, using the identical laser and detector settings for all animal cohorts. For 3D-reconstruction of dendritic, axonal, and mitochondrial and other organelle volumes, confocal stacks were deconvolved using the HuygensPro software (Scientific Volume Imaging, Netherlands) and further processed by the Imaris software (versions 9 and 10, Bitplane). 3D images were used to generate volume masks for cellular (using cytosolic reporters) and mitochondrial compartments, followed by calculation of mitochondrial parameters (volumes, occupancy etc.). Quantification of GABA and Vgat expression levels was performed by generating a volume mask of the axonal arbor using the tdTomato signal, which was then used to quantify colocalization over the thresholded signal values for both markers.

For MitoTimer analysis, raw dual-channel confocal z-stacks were split into green and red channel images using a custom ImageJ/Fiji macro. To enable unbiased segmentation independent of oxidation state, the two channels were summed pixel-wise using the ImageJ image calculator, and three-dimensional mitochondrial ROIs were defined on the composite images using the 3D ROI Manager plugin (3D ImageJ Suite) following conversion to 8-bit, with an intensity threshold of 30–255 applied uniformly across all images. All images were deconvolved (HuygensPro software, Scientific Volume Imaging, Netherlands) prior to segmentation to reduce out-of-focus background fluorescence. ROI sets were then reloaded onto the corresponding split green and red channel images, and the 3D Manager quantification function was used to extract mean fluorescence intensities from each channel for every ROI. For each mouse, at least 3 fields of acquisition per examined region were used for analysis, while identical imaging acquisition settings were maintained across groups. Downstream data processing was performed using custom Python scripts (Python 3; pandas, seaborn, and matplotlib libraries) in Jupyter Notebook via Anaconda. Per-mitochondrion green and red mean intensities were paired by ROI identifier. The green-to-red (G/R) fluorescence ratio was calculated for each mitochondrion and used to classify mitochondria as high-green (G/R > 1.2) or high-red (G/R < 0.8), whereas intermediate values (0.8 ≤ G/R ≤ 1.2) were excluded to reduce misclassification from overlapping central ROI values.

### Quantification and statistical analysis

Data are represented as mean ± standard error of the mean (SEM). Graphical illustrations and significance were obtained with GraphPad Prism (Version 8) or with OriginPro (OriginLab). Data in all groups were initially tested for normality of residuals via D’Agostino-Pearson omnibus and Shapiro-Wilk normality tests, followed by appropriate parametric or non-parametric analyses. Significance was defined as: ^∗^p ≤0.05, ^∗∗^p ≤ 0.01, ^∗∗∗^p ≤ 0.001. Group sizes (n, representing the number of samples or mice used) are indicated in the figure legends.

