## Supplementary material for "Mitochondrial turnover at central GABAergic synapses governs vulnerability to epileptic seizures": Key resource table

| REAGENT OR RESOURCE | SOURCE | IDENTIFIER |
| --- | --- | --- |
| Antibodies | | |
| Rabbit polyclonal anti-RFP | Rockland | Cat# 600-401-379; RRID: AB_2209751 |
| Rabbit polyclonal anti-Iba1 | Wako | Cat# 019-19741; RRID: AB_839504 |
| Chick polyclonal anti-GFP | Aves Labs | Cat# GFP-1020; RRID: AB_10000240 |
| Mouse monoclonal anti-GFAP, clone GA5 | Millipore | Cat# MAB360; RRID: AB_11212597 |
| rat monoclonal anti-Somatostatin | Millipore | Cat# MAB354; RRID: AB_2255365 |
| Guinea pig polyclonal anti-NeuN | Millipore | Cat# ABN90P; RRID: AB_2341095 |
| Mouse monoclonal anti-c-fos | Abcam | Cat# ab7963; RRID: AB_306177 |
| Rabbit polyclonal anti-Cre recombinase | BioLegend | Cat# 908001; RRID: AB_2565079 |
| Guinea pig polyclonal anti-Parvalbumin | Synaptic Systems | Cat# 195004; RRID: AB_2156476 |
| Rabbit anti-RHOT1 | Sigma | Cat# HPA010687; RRID: AB_1079813 |
| Goat polyclonal anti-Hsp60 | Santa Cruz | Cat# sc-1052; RRID: AB_631683 |
| Guinea pig polyclonal anti-b-3-TUBULIN | Synaptic Systems | Cat# 195004; RRID: AB_10805138 |
| Rabbit anti-Trak1 | Sigma | Cat# HPA005853; RRID: AB_1854778 |
| Rabbit anti-Trak2 | Sigma | Cat# HPA015827; RRID: AB_1858238 |
| Mouse monoclonal anti-b-Actin | Sigma | Cat# A2228; RRID: AB_476697 |
| Rabbit monoclonal anti-Snph | Abcam | Cat# ab192605; RRID: N/A |
| Rabbit monoclonal anti-Tom20 | Santa Cruz | Cat# SC-11415; RRID: AB_2207533 |
| Rabbit polyclonal anti-Rhot1 | Atlas antibodies | Cat# SC-11415; RRID: AB_1079813 |
| Guinea pig polyclonal anti-Vgat | Synaptic Systems | Cat# 131004; RRID: AB_887873 |
| Mouse monoclonal anti-Mtco1 | Abcam | Cat# AB14705; RRID: AB_2084810 |
| Mouse monoclonal anti-Atp5b | Abcam | Cat# AB5432; RRID: AB_304883 |
| Rat monoclonal anti-Lamp2 | Abcam | Cat# ab13524; RRID: AB_2134736 |
| Donkey anti-chicken IgY (H+L), Alexa Fluor 488 | Jackson ImmunoResearch | Cat# 703-545-155; RRID: AB_2340375 |
| Donkey anti-chicken IgY (H+L), Alexa Fluor 647 | Jackson ImmunoResearch | Cat# 703-605-155; RRID: AB_2340379 |
| Donkey anti-rabbit IgY (H+L), Alexa Fluor 647 | Jackson ImmunoResearch | Cat# 711-605-152; RRID: AB_2492288 |
| Donkey anti-goat IgY (H+L), Alexa Fluor 647 | Jackson ImmunoResearch | Cat# 705-606-147; RRID: AB_2340438 |
| Donkey anti-mouse IgY (H+L), Alexa Fluor 647 | Jackson ImmunoResearch | Cat# 715-605-150; RRID: AB_2340862 |
| Donkey anti-Rabbit IgG (H+L), Alexa Fluor 546 | Thermo Fisher Scientific | Cat# A10040; RRID: AB_2534016 |
| Donkey anti-Rat IgG (H+L) Alexa Fluor 647 | Jackson ImmunoResearch | Cat# 712-605-150; RRID: AB_2340693 |
| Donkey anti-Guinea Pig IgG (H+L) Alexa Fluor 647 | Jackson ImmunoResearch | Cat# 706-605-148; RRID: AB_2340476 |
| Rabbit anti-Mouse IgG-HRP | Sigma | Cat# A9044; RRID: AB_258431 |
| Goat anti-Rabbit IgG-HRP | Sigma | Cat# A0545; RRID: AB_257896 |
| Bovine anti-Goat IgG-HRP | Santa Cruz | Cat# sc-2384; RRID: AB_634814 |
| Chemicals, Peptides, and Recombinant Proteins | | |
| 4’,6-Diamidino-2-Phenylindole, Dihydrochloride (DAPI) | Thermo Fisher Scientific | Cat# D1306; RRID: AB_2629482 |
| Doxycycline hyclate | Sigma | Cat# D9891; RRID: N/A |
| Paraformaldehyde | Sigma-Aldrich | Cat# P6148; RRID: N/A |
| Tetramethylrhodamine methyl ester (TMRM) | Thermo Fisher Scientific | Cat# T668; RRID: N/A |
| Critical commercial assays | | |
| Adult Brain Dissociation Kit | Miltenyi Biotec | Cat# 130-107-677 |
| Bacterial and Virus Strains | | |
| AAV1-FLEX-mito-  GFP.WPRE-SV40 | Benoit et al., 2025 | N/A |
| AAV1-FLEX-EGFP2-  Gephyrin P1 | This paper | Modified from Addgene, Cat# 68815; RRID:Addgene_68815 |
| AAV-EF1a-DIO-PSD95-  EGFP-WPRE | Sun et al., 2019 | Addgene, Cat# 133785, a gift from Michael Stryker; RRID: Addgene_133785 |
| AAV-DIO-LACTB-dAPEX2 | Zhang et al., 2019 | Addgene, Cat#  117180, a  gift from David  Ginty; RRID:Addgene_117180 |
| AAV-CAG-DIOerHRP(N175S mutant)-  WPRE.SV40 | Joesch et al., 2016 | Addgene, Cat# 79906, a  gift from Joshua  Sanes; RRID:Addgene_79906 |
| AAV1-CAG-GFP-WPRE.SV40 | This paper | N/A |
| AAV1-CAG-GFP-T2A-iCre | This paper | N/A |
| AAV1.CamKII 0.4-Cre.SV40 | This paper | N/A |
| AAV-1-FLEX-Miro1-T2A-mito-  GFP-WPRE.SV40 | This paper | N/A |
| AAV-1-FLEX-SNPH-T2A-mtGFP | This paper | N/A |
| AAV1 -FLEX-pTRE-Tight-MitoTimer | This paper | Modified from Addgene, Cat# 50547, a gift from Roberta Gottlieb; RRID:Addgene_50547 |
| AAV1-FLEX-rtTA | This paper | N/A |
| AAV-EF1a-double floxed-hChR2(H134R)-EYFP-WPRE-HGHpA | Addgene | Addgene, Cat# 20298, a  gift from Karl Deisseroth; RRID:Addgene_20298 |
| AAV1-FLEX-mtTurquoise2 | This paper | N/A |
| AAV1-FLEX-emerald-Lamp1 | This paper | N/A |
| AAV-PHP.eB-FLEX-mtGFP | This paper | N/A |
| AAV-PHP.eB-FLEX-Miro1-T2A-mtGFP | This paper | N/A |
| AAV-PHP.eB-FLEX-Trak1-myc-DDK | This paper | N/A |
| Experimental Models: Organisms/Strains | | |
| Mouse: Thy1^GFP^ | Feng et al., 2000 | Tg(Thy1-EGFP)MJrs; RRID: N/A |
| Mouse: SST^CRE^ | Taniguchi et al., 2011 | B6N.Cg-*Sst^tm2.1(cre)Zjh^*/J; RRID:IMSR_JAX:018973 |
| Mouse: PV^CRE^ | Hippenmeyer et al., 2005 | B6.129P2-*Pvalb^tm1(cre)Arbr^*/J; RRID:IMSR_JAX:017320 |
| Mouse: tdTomato^LSL^ | Madisen et al., 2010 | B6.Cg-*Gt(ROSA)26Sor^tm14(CAG-tdTomato)Hze^*/J; RRID:IMSR_JAX:007914 |
| Mouse: Vgat^CRE^ | Vong et al., 2011 | Slc32a1^tm2(cre)Lowl^; RRID:N/A |
| Mouse: mtYFP^LSL^ | Sterky et al., 2011 | Gt(ROSA26)SorStop-mito-YFP; RRID: N/A |
| Mouse: *Miro1* floxed | Nguyen et al., 2014 | B6(Cg)-*Rhot1^tm2.1Jmsu^*/J; RRID: RRID:IMSR_JAX:031126 |
| Mouse: *Trak1* floxed | This paper | C57BL/6N-Trak1^em389Cecad^/Cecad; RRID: N/A |
| Mouse: *Trak2* floxed | This paper | C57BL/6N-Trak2^em362Cecad^/Cecad; RRID: N/A |
| Mouse: *Snph* flexed | This paper | C57BL/6N-Snph^em393Cecad^/Cecad; RRID: N/A |
| Software and Algorithms | | |
| ImageJ | NIH | <http://imagej.nih.gov/ij/>; RRID: SCR_003070 |
| Fiji | Max-Planck-Gesellschaft | <http://fiji.sc>; RRID: SCR_002285 |
| Photoshop | Adobe | https://www.adobe.com/; RRID: SCR_014199 |
| GraphPad Prism | GraphPad Software | https://www.graphpad.com/scientific-software/prism; RRID: SCR_002798 |
| Huygens Professional | Scientific Volume Imaging | RRID: SCR_014237 |
| Adobe InDesign | Adobe | https://www.adobe.com/; RRID: CR_021799 |
| Imaris | Bitplane | http://www.bitplane.com/imaris/imaris; RRID:SCR_007370 |
| Signal | Cambridge Electronic | http://ced.co.uk/products/signal; RRID:SCR_014276 |
| IgorPro | WaveMetrics | http://www.wavemetrics.com/products/igorpro/igorpro.htm; RRID:SCR_000325 |
