## Supplemental table 1 for "Mitochondrial turnover at central GABAergic synapses governs vulnerability to epileptic seizures"

### Supplemental Table 1: Sequences of Cas9 crRNA and DNA repair templates

| gRNA sequence (5'-3') | Target |
|---|---|
| CAGGTATTTCCGCTAAGGTG | Snph, intron 4 |
| GTGGGTTCTGACAAAGTGAG | Trak1, upstream of exon 5 |
| CTTTGCCCTGGCACATCCGT | Trak1, downstream of exon 5 |
| GAGTTTCTTCAGGTCTGGAC | Trak2, upstream of exon 4 |
| CTTCAGGGGCTCTTCTACTC | Trak2, downstream of exon 4 |

| DNA repair template sequence (5'-3') | Description |
|---|---|
| GAGACTGGGAGTCTCAAATGGGAGATGTGTTATAAA<br>ACTGTAAGTAACAGAAAAGTCTGGAAGCTTCTAATT<br>CCAGGCTTTGCAAGATGCAAATCAGAAGCCCGAGA<br>CTGGGCACGTGTCTTACACATGACTGTTACCTACCA<br>TCCACACATAACTTCGTATAGCATACATTATACGAAG<br>TTATTCTTTGCACCATTCTAAAGAATAACAGTGATAA<br>TTTCTGGGTTAAGGCAAATAACTTCGTATAGGATACT<br>TTATACGAAGTTATTAGGGCGCAGTAGTCCAGGGTT<br>TCCTTGATGATGTCATACTTATCCTGTCCCTTTTTTT<br>TCCACAGCTCGCGGTTGAGGACAAACTCTTCGCGG<br>TCTTTCCAGTCGACTGACTGACTGAATTCAATACACT<br>CTTCTTCAACGCCTGCTGCCTGGGCTTCGTTGCCTA<br>TGCCTACTCTGTGAAGGTGAGTGTGCTAGAGGGGG<br>TGGGAGGGAATGGTGTAAGAAGGTGTGTGTGAGCC<br>TGAAGCGCCATCTACTCAGATAGCACCGAGGTGTG<br>AGTGTGGGGTCCTTTCTGTAAGAAAGTGTGTGTGCA<br>CCCTTCCTATTTGTGTGTGTGTATAACTTCGTATAAT<br>GTATGCTATACGAAGTTATTAAGAGGTTTCATATTGC<br>TAATAGCAGCTACAATCCAGCTACCATTCTGCATAA<br>CTTCGTATAAAGTATCCTATACGAAGTTATCTTAGCG<br>GAAATACCTGAGATCTTTAGTGGCTCTGGGCATCCC<br>TGAACACAGATGATGTCTCTCTACCGAGGCGGAAG<br>AGCTTGACTGTGGGACTGCTGTCAGAACATACCTGT<br>CCTTGTGTTCATGACCCTCCAAGCCTCACTGTGGTC | Snph FLEx cassette (depicted 5'-3' on noncoding DNA strand; inverted features underlined)<br>homology_arm/loxP/linker/lox2272/Ad_splice_acceptor/3xStop/Ifitm2_splice_donor/loxP/linker/lox2272/homology_arm |
| ACAACACAGCTGTGTCCCTACAGCAGGCCACTAAG<br>CTCCCAAGAGCAAGGGTCTCCTGGCTTTCACAGAA<br>GCATCCAGACTTCTGTCCCAAGGACCTGAGGCTCC<br>GGGATCTATGTTTTCATCCACCATTGAGCACACCCC<br>ATTCCCACGATAACTTCGTATAATGTATGCTATACGA<br>AGTTATGATGTGCCAGGGCAAAGGGCCACTCTGAC<br>AAGCCAGGATTGACACCTTGTAAGAGAGAAACTGCC<br>CTCCACTCAGCAGCTCACTTGAAGCAGTGAGGTGG<br>TGTCATTTGAGGCCTTCAGATGGCCGTTCTGGGAAA<br>GACTATCCCCAAGACGCTAGTGGCCCTGCCTAGGG<br>GCCTCCAAGGGCACCCATTGCACATCAGCCCGCCT<br>GCCCTCAGCTCTGGGCACCTGTGCTGGAGACTTAC<br>GGGGTTGAGCAGACGGACTCAGGCTCGCTCTCCTC<br>AGCGGCACTGGTGTAGAATTGAAGCAGCTCGTCTTT<br>CATGGACAGCTCATGTCGGAGCTGAGACACCTGTG<br>CGGACAGCAAGGGGGCAGATCGATGCTTGCTGGAA<br>GGTGGCAACGTCTTTCTGTGCTCGAACGCCAGAGA<br>GGGGTGGCATCCCAAGTGAAGGCTGGTTTGCTACT<br>CAGAGTCTCAAAGCCTTGAGTTTGACAATGTTTAAG<br>AAAAGCCCAGTCCATCAGTTCTGGGTTGGAAACGGT<br>CTATGTCCTCACATTTGGTCAGAAGAGCAGAGAGG<br>GGAGCAAGCAGAAGGCTCTGGAATGATCAGGGGGT<br>TGATGCTTGTTGGCTCTGGAAGCAGCCAGGTGACC<br>AGCTGGGCAAGTGGCCAGAACAGTGGGCCATGATA<br>CTAAATAAAATGAGAGGCCACTCGAATTCATAACTT<br>CGTATAATGTATGCTATACGAAGTTATACTTTGTCAG<br>AACCCACCCAGTGCCATCTCAGAGCAGATCTATGG | Trak1, loxP flanked exon 5 (depicted 5'-3' on noncoding DNA strand)<br>homology_arm/loxP/intron /exon/intron/EcoRI/loxP/homology_arm |

<table>
<tr><td>CTGTGCTGCCTCTGGGTGGGCTGCCAGACTCCACC<br>CCTCTTAGAAAACCACACAGCCCTCCATGTGACCAG<br>CACACCTCAGTGTCCCTCAACAAGCTGGTCAGAC</td><td></td></tr>
<tr><td>GTGATCCTGGGACCCTGAGATCCTGGTGTGACCAA<br>GCTCCTGGGATCCTGGGATCCTGGGCGTAGTAGAG<br>AGCCTGGGAATGGAGCTTCCTCTGGGTATTGTGGG<br>ACTGGCTGTGGAGTTTGAGCCCAAGGTCTGCTCAG<br>GGCACCGGTCATAACTTCGTATAGCATACATTATAC<br>GAAGTTATGAATTCCAGACCTGAAGAAACTCGTGCT<br>ACTGGTTGGGCGGTGTTCCTGGGTGCCTGGGTTCC<br>TCTGGTCCTAGTTACTTCCTAAAGTAACTTGTTTTTA<br>ACTGTCTTAGTTCTAGGCACAGACAGAGTGGAGCA<br>GATGACCAAAACCTACAATGACATTGACATGGTCAC<br>ACATCTCCTGGCAGAGGTAGGCATGTATCTTTTCTC<br>TCTCTGGTATAAAACCTAAAGCTTGAGGGTCGTTTG<br>GAATGTGCCGGGTTTGCAGAGATAAAACATGATGCT<br>CAGCCTGTTGTCTAGACTGTTTTGCATTGATTGTTAT<br>AAGTGATATACTAGCTGGCATCCTGAGATAACTTCG<br>TATAGCATACATTATACGAAGTTATTAGAAGAGCCC<br>CTGAAGCTCATGATTTTGATGTGGATGGATTGGCAT<br>ATATAGTGTCGTCTGACCTCCCTGTCTTAGGATAAA<br>GCAATTGTTTAAAAAGCCTTTATTTTTAACTATGGGG<br>GACAATGCTCTTGGGCACCGAAGGCTGTGA</td><td>Trak2, loxP flanked exon 4 (depicted 5’-3’ on coding DNA strand)<br>homology_arm/loxP/EcoRI/intron /exon/intron/ loxP/homology_arm</td></tr>
</table>
